# Heterologous reporter expression in the planarian *Schmidtea mediterranea* through somatic mRNA transfection

**DOI:** 10.1101/2021.04.20.440701

**Authors:** Richard Nelson Hall, Uri Weill, Leonard Drees, Sergio Leal-Ortiz, Hongquan Li, Chew Chai, Alejandro Sánchez Alvarado, Nicholas A. Melosh, Andrew Z. Fire, Jochen C. Rink, Bo Wang

## Abstract

Planarians have long been studied for their regenerative abilities. Moving forward, tools for ectopic expression of non-native proteins will be of substantial value. Using a luminescent reporter to overcome the strong autofluorescence background of planarian tissues, we demonstrate heterologous protein expression in planarian cells and live animals. Our approach is based on the introduction of mRNA through several nanotechnological and chemical transfection methods. We improve reporter expression by altering untranslated region (UTR) sequences and codon bias, facilitating measurement of expression kinetics both in isolated cells and in whole planarians using luminescence imaging. We also examine protein expression as a function of variations in the UTRs of delivered mRNA, demonstrating a framework to investigate gene regulation at the post-transcriptional level. Together, these advances expand the toolbox for the mechanistic analysis of planarian biology and establish a strong foundation for the development and expansion of transgenic techniques in this unique model system.

**Motivation:** The study of planarians has contributed to advances in our understanding of regeneration, stem cell dynamics, and many other fundamental biological processes. However, the persistent challenge of expressing transgenes in planarians has led to the speculation that they may be resistant to transfection. In this work, we develop methods to express exogenous mRNAs in both isolated planarian cells and whole animals by optimizing delivery techniques, genetic constructs, and detection methods. These methods allow us to study transfection kinetics and post-transcriptional regulation of gene expression in a quantitative manner. Beyond planarian research, this work should also provide a broadly applicable strategy to develop similar tools for animals that are also challenging to modify genetically.

## Introduction

Planarian flatworms have fascinated generations of scientists with their regenerative abilities and have played a critical role in our efforts to understand stem cells and regeneration (Newmark & Sánchez Alvarado, 2002; Reddien, 2018; Rink, 2018). Planarians can regenerate their entire body from a small tissue fragment. During this process, they restore their body axes and rebuild all organs to have appropriate proportions. While gene knockdown by RNA mediated genetic interference (RNAi) (Sánchez Alvarado & Newmark, 1999; Reddien et al., 2005; Collins et al., 2010) and next generation sequencing techniques (Böser et al., 2013; Lakshmanan et al., 2016; Guo et al., 2017; Grohme et al., 2018; Fincher et al., 2018; Plass et al. 2018; Guo et al., 2021) have been widely used in planarian research, tools for transgene expression are still needed. Earlier efforts have attempted whole animal electroporation of plasmid DNA encoding a fluorescence protein (González-Estévez et al., 2003), but the intense autofluorescence of planarian tissues and lack of orthogonal verification of transgene expression have limited the applicability of this study.

In general, establishing reporter expression in a system needs to overcome three challenges: construct delivery, expression, and detection. Delivery and expression are necessary prerequisites for detection, while, on the other hand, delivery and expression need reliable detection to optimize. In addition, strategies to circumvent various genetic defense mechanisms (Kim et al., 2019; Chen et al., 2003; Aljohani et al., 2020) that may restrict transgene expression are difficult to test without a robust reporter. Therefore, the first demonstration of reporter expression is often the bottleneck for developing transgenic tools; establishing a positive control can transform the method development process from a random walk in a vast parameter space into a well constrained optimization problem.

Injecting constructs into embryos often provides the first route for transgene expression in most systems. However, it is infeasible to inject in planarian’s ectolethical eggs, as blastomeres are tiny and dispersed among yolk cells (Cardona et al., 2006; Davies et al., 2017). Furthermore, the commonly used planarian strains of *Schmidtea mediterranea* are exclusively asexual, reproducing through fission and regeneration (Benazzi et al., 1972; Vila-Farré & Rink, 2018). This leaves introduction of genetic material into individual somatic cells as the most general approach to genetic modification in this system. While possible in some vertebrate models and immortalized cell lines, transformation protocols and reagents are typically optimized for specific systems. In addition, measuring delivery efficiency relies on expression as the ultimate readout. Due to the distant relationship between planarians and other model organisms, it is unclear what modifications to reporter constructs are needed to drive expression in planarians. Finally, autofluorescence limits the utility of fluorescent proteins in planarians, especially during the initial stages of reporter optimization when signals may be weak.

Here, we report a robust and extensively verified method for heterologous gene expression in planarians. To address the problem of delivery, we used a direct nanoscale injection method to first establish a positive control, based on which we identified efficacious chemical transfection reagents for transforming planarian cells both *in vitro* and *in vivo*. To circumvent the numerous variables associated with DNA transfections, we delivered synthetic mRNA for ectopic expression. Finally, for detection, we relied on luminescent reporters, in particular, a compact, bright, and stable luciferase, nanoluciferase, Nluc (Hall et al., 2012; England et al., 2016). Using this sensitive and quantitative luminescence readout, we improved the reporter construct by altering untranslated regions and codon usage biases and presented a case study in which we identified regulatory sequences that modulate gene expression post-transcriptionally. Via luminescence imaging, we quantified single-cell transfection kinetics and explored limiting factors in transgene expression in planarian cells. Finally, we demonstrated the utility of luminescence imaging for monitoring gene expression in live animals. Our results not only provide the first positive control for exogenous gene expression in planarians to guide the future development of planarian transgenesis, but also offer a new route to measure and understand gene expression and regulation in planarian cells.

## Results

### Nanoluciferase mRNA delivered through nanostraws is expressed in planarian cells

To establish mRNA expression in the asexual strain of *S. mediterranea*, we sought to identify an efficient platform for delivering genetic material into primary planarian cells. We selected nanostraws, which combine the robustness of microinjection with the throughput of bulk electroporation (Tay & Melosh, 2019).

Nanostraws are approximately 100-200 nm wide hollow aluminum oxide tubes protruding from a polycarbonate substrate situated above a buffer reservoir containing the genetic material to be delivered (**Figure 1A-B**). Cells are centrifuged against the straws which engage in close contact with the cell membrane. Electric pulses are then used to locally porate the membranes and electrophorese genetic material through the nanostraws and directly into the cellular cytoplasm (**Figure S1**). By only permeabilizing membranes immediately in contact with nanostraws, they have been shown to improve both delivery efficiency and viability of transformed cells (Xie et al., 2013; Cao et al., 2018).

**Figure 1:**
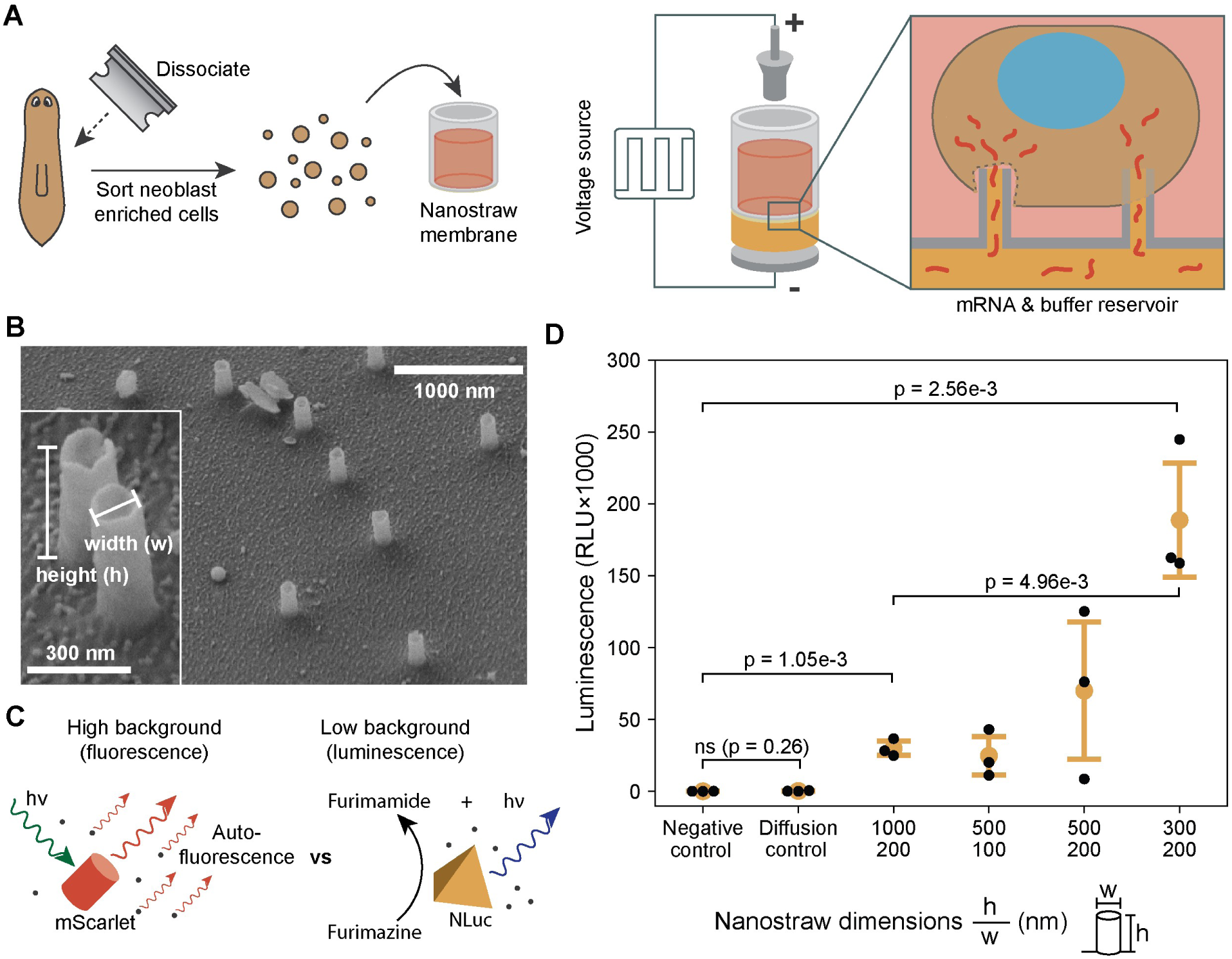
Nluc mRNA delivered through nanostraws is expressed in planarian cells. **(A)** Schematics showing the steps of nanostraw delivery. In each experiment, 200,000 cells are placed into a nanostraw cartridge and electroporated with square wave pulses between two titanium electrodes. **(B)** SEM images of nanostraws. Inset: a magnified view showing individual straws. **(C)** Unlike autofluorescence, autoluminescence is absent in planarian tissues, making luminescent reporters easy to detect. **(D)** Luminescence from 200,000 cells at 24 hpt with sNluc1 mRNA. Cells are transfected using three 45 s square wave pulse train of 35 V, 200 μs, and 40 Hz with 1 min between each cycle. Negative control: nanostraw delivery with PBS alone. Diffusion: mock experiments without applying electrical pulses. Luminescence is measured at room temperature with NanoGlo-Live substrate by plate reader. Each data point represents one biological replicate which was conducted on an independently isolated population of cells using nanostraws from the same batch and mRNA synthesized in the same reaction. Error bars: standard deviation. Statistical analysis is performed with one-way ANOVA (p = 8.05e-5) in addition to two-sided Welch’s t-test to calculate p-values between pairs of conditions; ns: not significant (p > 0.05).

To prepare cells for nanostraw delivery, we flow-sorted a stem cell (i.e., neoblast) enriched population based on their light scattering properties, so-called X1FS, from dissociated planarians (Hayashi et al., 2006; Wagner et al., 2011). This population is uniform in size and depleted of debris, which are important preconditions for nanostraw delivery. Our initial experiments used capped and polyadenylated *in vitro* transcribed mRNA encoding the red fluorescent protein, mScarlet, fused to the planarian histone H2B (**Table S1**). We reasoned that red fluorescence might be more easily detectable against planarian autofluorescence (Lim et al., 2019), which is biased towards shorter wavelengths, and that a nuclear localization signal might further enhance signal-to-noise. We used a pulse train protocol that was empirically optimized for delivery in human hematopoietic stem cells, which have similar morphological characteristics with XIFS cells, i.e., small size, minimally adherent, high nucleus to cytoplasm ratio (Schmiderer et al., 2020).

We performed flow cytometry to quantify fluorescence signal and found, even in this relatively uniform cell population, a broad distribution of fluorescence intensities spanning three orders of magnitude in both experimental and negative conditions (**Figure S2**), making the assessment of transfection outcomes difficult. Since some cells genuinely exhibit brighter fluorescence than others, false positives may be common and true positives could be obscured by the broad autofluorescent background. These results compelled us to seek an alternative non-fluorescent reporter to unambiguously quantify gene expression in planarian cells.

Unlike fluorophores, luciferases produce light as the result of an oxidative chemical reaction with an exogenously added substrate (luciferin), and most animal tissues are devoid of autoluminescence due to the enzymatic nature of luciferases (**Figure 1C**). Therefore, we delivered mRNA encoding a planarian codon optimized nanoluciferase (sNluc1) (**Table S1**) into X1FS cells. Transfected cells were maintained for 24 hr in Iso-L15, a nutrient rich medium with reduced osmolarity. A similar medium has been shown to improve planarian cell viability (Lei et al., 2019). We observed a clear and reproducible luminescence signal in transfected cells over 100-fold above the background signal of the negative controls. The physical dimensions of the nanostraws strongly influenced the signal intensity, as has been observed in previous nanostraw transfection experiments (Xie et al., 2013), implying that we were in a regime where expression was dependent on the amount of mRNA delivered (**Figure 1D**). Overall, these results provide a proof-of-principle for heterologous protein expression in planarian cells.

### A screen identifies chemical reagents to efficiently transfect planarian cells

With a validated reporter construct in hand, we next sought to identify a more broadly accessible mRNA delivery method. Chemical transfection reagents have become increasingly popular because of their ease of use, high efficiency, and scalability, but they are often developed and optimized for specific cell types.

To identify suitable reagents for planarian cell transfection, we screened a panel of commercially available reagents by transfecting *in vitro* transcribed sNluc1 mRNA containing 5’ and 3’ UTR sequences from a highly expressed planarian gene, Y-box binding protein (*YB1*) (**Table S1**), and measuring the luminescence on a plate reader. Our screen used total cells from freshly dissociated animals rather than X1FS cells due to the large number of transfections needed (**Figure 2A**). Initially, all reagents were utilized according to the manufacturer’s recommended protocols (**Table S2**). While most transfection reagents produced little to no expression, Viromer and Mirus Trans-IT reagents, both of which are comprised of endoosmolytic cationic polymers, consistently achieved luminescent signals 100 to 1,000-fold above the negative control (**Figure 2B**).

**Figure 2:**
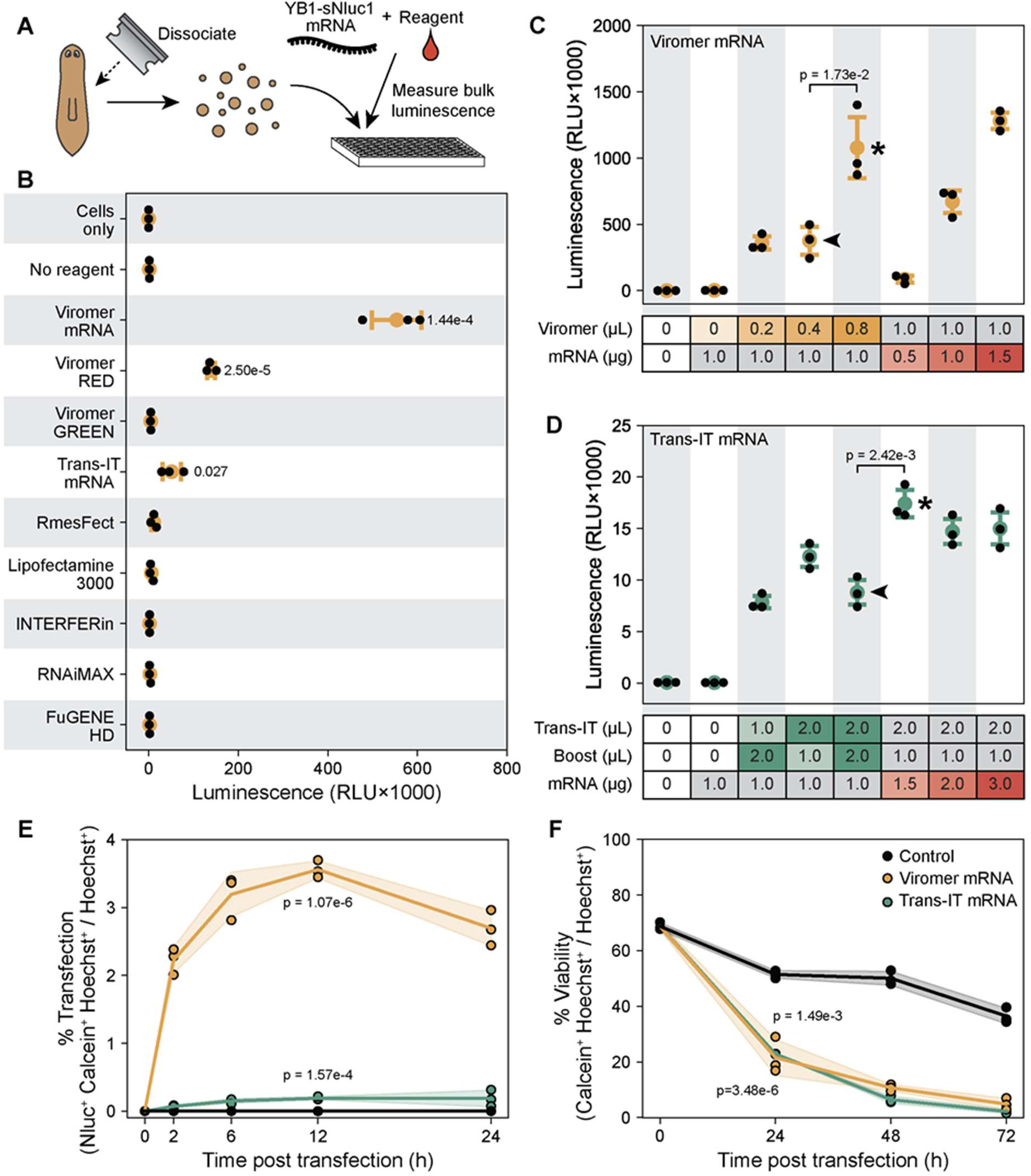
A screen identifies chemical reagents to efficiently transfect planarian cells. **(A)** A diagram demonstrating the chemical transfection reagent screen pipeline. **(B)** Luminescence from 200,000 cells transfected with 1 µg of YB1-sNluc1 mRNA delivered by chemical transfection reagents, using the manufacturer’s recommended protocols, assayed at 24 hpt (**Table S2**). (**C-D**) Optimization of Viromer mRNA (**C**) and Trans-IT mRNA (**D**) transfections identifies conditions with increased expression that are used for all following experiments (asterisk), compared to the condition initially used in the screen (arrowheads). 200,000 cells are transfected in each experiment and assayed at 24 hpt. **(E)** Percentage of luminescent cells (Nluc^+^Calcein^+^Hoechst^+^ / Hoechst^+^) over the course of 24 hpt. **(F)** Cellular viability (Calcein^+^Hoechst^+^ / Hoechst^+^) over the course of 72 hpt. Cells in (**E**) and (**F**) are transfected according to the optimal conditions found in (**C**) and (**D**) with either Viromer (yellow), Trans-IT (green), or left untransfected (black). Cells are imaged on an Olympus LV200 luminescence microscope using a 20× air objective (Olympus, UPLXAPO20X). Cells are stained with Hoechst 33342 (1 μM) and Calcein AM (1 μg/μL) and quantified using scanR Analysis 3.2.0. Luminescence is measured at room temperature with NanoGlo-Live substrate. Statistics. Data points in (**B**) represent the mean of technical replicates (n = 3) for each biological replicate. Data points in (**C**-**F**) represent technical replicates using mRNA prepared from the same batch. Error bars: standard deviation. ANOVA: p = 2.08e-18 (**B**); p = 1.25e-9 (**C**); p = 2.05e-11 (**D**). p-values for pairwise comparison are calculated using two-sided Welch’s t-test and reported in the figure. In (**B, E, F**), comparisons are made by comparing experimental groups with the negative control in which mRNA is delivered with no transfection reagent.

To further optimize the transfection protocols, we tested various ratios between Nluc mRNA and reagent components (**Figure 2C-D**). For Viromer, increasing both the amount of reagent and/or mRNA in the transfection mix doubled the baseline signal, with 0.8 µL of reagent per 1 µg mRNA per reaction as an economic compromise for further experiments. A similar two-fold boost over the baseline signal was achieved for Trans-IT, but maximal signal intensities in cell cultures remained below those achieved by Viromer transfections. These conditions, however, still only produced luminescence ∼100-fold lower compared to transfections of mammalian cells (e.g., HeLa cells) (**Figure S3**), indicating that future optimizations might help to achieve substantially higher expression levels.

To confirm that luminescence was produced by transfected planarian cells, we imaged cells transfected with nanostraws, Trans-IT, and Viromer using NanoGlo-Live furimazine substrate (Promega) on an LV200 bioluminescence imaging system (Olympus). Individual luminescent cells were evident in transfected samples (**Figure S4**) but never observed in negative controls. High magnification imaging confirmed the cytoplasmic origin of the luminescence signal. By counting luminescent live cells over time, we quantified the transfection efficiency to be ∼3.5% out of all cells for Viromer, which plateaued starting ∼12 hr post transfection (hpt), and 0.2% for Trans-IT (**Figure 2E**), whereas cell viability was similar after both transfections (**Figure 2F**).

Comparing both reagents, the discrepancy in percentage of luminescent cells is smaller than the difference in total luminescence intensity measured by the plate reader, suggesting that Viromer transfected more cells and produced higher expression per cell. Finally, to further demonstrate that Nluc mRNA is the source of the luminescence, we created a Nluc variant containing a premature stop codon, which completely abolished expression and thus provided additional evidence for transgene expression in planarian cells (**Figure S5**). Overall, these observations establish chemical delivery as a viable method for mRNA transfection of planarian cells, with Viromer and Trans-IT as promising leads.

### Altering untranslated region (UTR) sequences and codon bias improves Nluc expression

With efficient delivery reagents, we sought to improve the reporter construct. To identify UTRs that may enhance expression, we flanked sNluc1 with UTRs (**Figure S6-7**) from four endogenous genes with high expression in all cell types, reasoning that they may enhance Nluc mRNA stability and/or translation efficiency (see **Table S3** for sequence details). These UTRs either increased (RPL15, YB1, RPL10) or decreased (ENO) the expression of Nluc relative to the construct lacking endogenous UTRs (**Figure 3A**).

**Figure 3:**
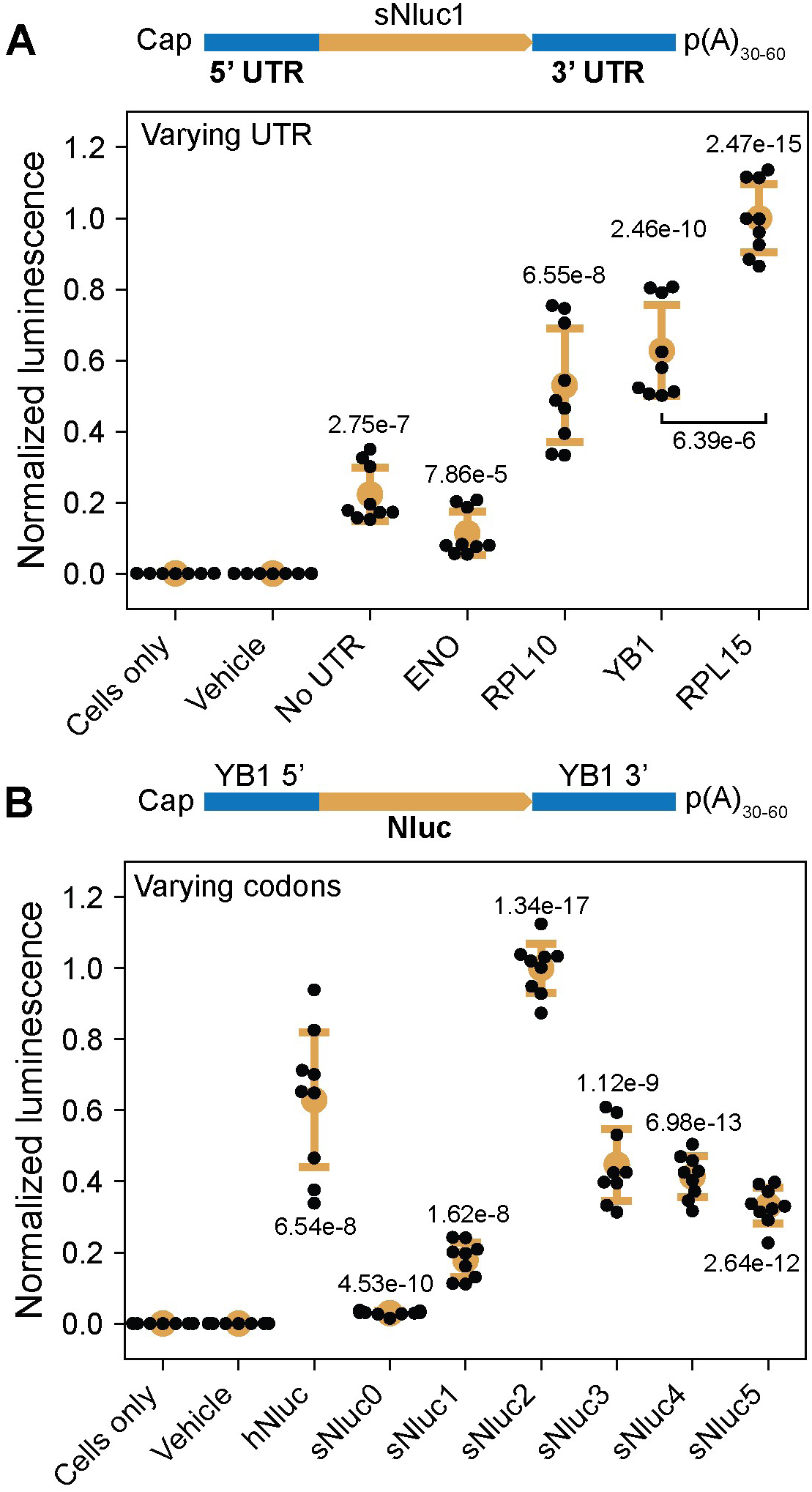
Improving Nluc construct by altering UTR and codon bias. **(A)** Luminescence from transfections using constructs incorporating different endogenous 5’ and 3’ UTRs flanking sNluc1. No-UTR mRNA contains flanking attB1 and attP1 sequences used for cloning. UTRs were cloned from *ENO*, enolase (dd_Smed_v6_510_0_1); *RPL10*, ribosomal protein L10 (dd_Smed_v6_130_0_1); *YB1*, Y-box binding protein (dd_Smed_v6_52_0_1); *RPL15*, ribosomal protein L15 (dd_Smed_v6_193_0_1), which are identified using the gene annotation provided by PlanMine (Rozanski et al., 2019). We also choose to use UTRs from shorter genes which mapped uniquely to the genome, allowing them to be more easily isolated via PCR. All transfections in this figure are performed with 200,000 cells from whole dissociated planarians using 0.8 µL Viromer and 1 µg mRNA per well and assayed 24 hpt at room temperature using NanoGlo-Live substrate. **(B)** Luminescence from alternative codon usage variants of Nluc flanked by YB1 UTRs. Codon optimization schemes tested include utilizing only the most frequent codon for each amino acid (sNluc0), the IDT online codon optimization tool (sNluc5) with manual adjustments to codon usage (sNluc1), and random sampling of the codon table biased by the frequency of each codon (sNluc2-4). Statistics. All data points presented are technical replicates (n = 3) from independent biological replicates (n = 3) each utilizing mRNA produced from independent *in vitro* synthesis reactions. Luminescence measurements for each panel are normalized by the mean of the highest intensity construct (panel **A**: RPL15; **B**: sNluc2). Error bars: standard deviation. ANOVA: p = 4.05e-31 (**A**); p = 2.98e-40 (**B**). p-values reported in the figure are calculated using two-sided Welch’s t-test between experimental groups and negative controls with cells only unless specified otherwise in the figure.

Next, we investigated the effect of codon optimization, the commonplace practice of matching endogenous codon biases in order to maximize expression (Quax et al., 2015; Jeacock et al., 2018). This may be important for *S. mediterranea*, in which a strong preference for A/T at third-base positions contributes to a genome-wide A/T-bias of 70% (Grohme et al., 2018). Therefore, we generated five additional codon-optimized Nluc variants, besides sNluc1, to cover a range of codon adaptation indices (CAI), a 0-1 bounded value measuring the gene’s codon bias compared to a reference codon preference table (Sharp & Li, 1987; Puigbò et al., 2008). We also included a Nluc sequence optimized for mammalian expression (hNluc) as a comparison. Unexpectedly, the luminescence in transfected cells did not correlate with CAI (**Figure 3B**). The most highly expressed construct (sNluc2, CAI = 0.713) was part of a series of constructs with very similar CAIs and GC content generated by randomly sampling the planarian codon table (sNluc2-4; **Tables S1, S3**). Based on these results, we combined RPL15 UTR and sNluc2 into an improved reporter construct that was used for all subsequent experiments unless otherwise explicitly specified (**Figure S8**).

### Live luminescence imaging reveals expression kinetics in vitro

Quantifying expression levels across cells and changes within single cells can help reveal important kinetics of transfection and gene expression. For quantitative microscopy, we built a customized luminescence microscope (Kim et al., 2017). Using a demagnifying tube lens, we expanded the field-of-view (FOV) of high magnification/high numerical aperture (NA) objectives that are needed for efficient light collection. In addition, we used a back-illuminated EMCCD camera (Andor iXon) for detection, which has single-photon sensitivity and provides quantitative measurements. The conventional substrate for Nluc, furimazine (Fz), has poor water solubility and is supplied in organic solvents which may stress cells and affect gene expression kinetics. To overcome this, we utilized a novel Fz derivative, fluorofurimazine (FFz), which has improved water solubility and thereby reduces cytotoxicity while maintaining Nluc’s characteristic brightness (Su et al., 2020).

With these technical advances, we quantified the distribution of luminescence intensities across transfected cells. For this experiment, bulk dissociated cells were allowed to adhere to a glass-bottom imaging well treated with concanavalin A to prevent them from moving in and out of the imaging plane. We first imaged cells at 24 hpt, as at this time point the number of luminescent cells should have plateaued (**Figure 2E**). In a single FOV (∼1.6 mm^2^), we captured over 80 individual luminescing cells (**Figure 4A**). From these images, we segmented cells and quantified their luminescence at 40 min post FFz addition, to allow sufficient time for FFz to diffuse into the cells. This revealed a long-tailed distribution with a small number of intensely bright cells (**Figure 4B**). Unexpectedly, we captured new transfection events even at this late time point, indicating that not only were these cells healthy enough to be transfected, but also that there remained functional Viromer-mRNA complexes in the media (**Figure 4C**).

**Figure 4:**
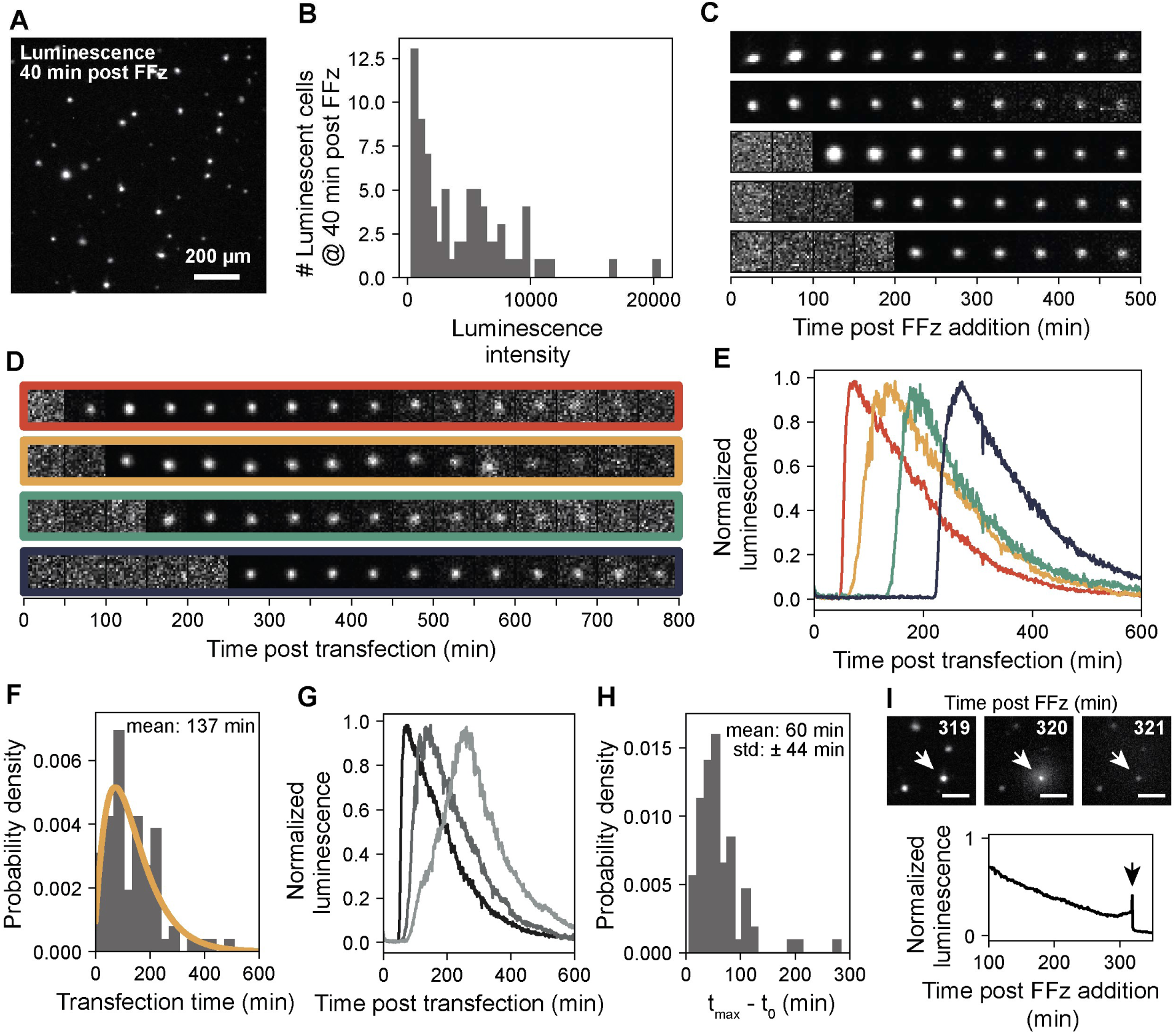
Time-lapse luminescence imaging reveals expression kinetics in single cells. **(A)** Representative luminescence image of transfected bulk dissociated planarian cells. FFz (1:250) is added at 24 hpt and the image is acquired 40 min post FFz addition. All transfections reported in this figure use 0.8 µL of Viromer mRNA and 1 µg of RPL15-sNluc2 mRNA per 200,000 cells. All luminescence imaging is performed at room temperature with FFz substrate. **(B)** The distribution of luminescence intensities of single cells treated as described in (**A**). Time of transfection is defined as the first time point when the total luminescence intensity of a cell exceeds ∼3-fold above the background intensity. **(C)** Examples showing temporal evolution of single-cell luminescence starting at 24 hpt when FFz is added. The three cells at the bottom are new transfections during imaging. **(D)** Representative images showing luminescence intensity changes in individual cells. FFz substrate and transfection complexes are added simultaneously at time 0, when imaging began immediately. **(E)** Time traces of single-cell luminescence intensities normalized against the maximum intensity of each cell. Luminescence traces correspond to the cell bound by a frame of the same color in (**D**). **(F)** Distribution of transfection times. The curve is fit for a gamma distribution (α = 2.3, β = 64.6). **(G)** Example luminescence traces showing individual cells increasing in luminescence at different rates. **(H)** Distribution of the time intervals between the time of transfection (t_0_) and the time of maximum luminescence (t_max_). **(I)** (Upper) Images of a transfected cell undergoing cell death and releasing Nluc at 319 mpt (immediately before rupture), 320 mpt (at the moment of rupture), and 321 mpt (after rupture). Scale bars, 100 µm. (Lower) Normalized luminescence intensity of the cell shown above. The arrow highlights the moment of rupture at 320 mpt.

Motivated by the observation that we could capture individual transfection events, we performed imaging from the moment of transfection to measure the kinetics of transfection and the onset of expression. We added transfection complexes and FFz at the same time and immediately began imaging. The sample started completely dark, but within the first hour many cells began to luminesce (**Figure 4D**, **Video S1**). We quantified the luminescence of each cell (**Figure 4E**) and extracted the transfection time, defined by the time when the luminescence exceeds the background noise. The transfection time followed a gamma distribution (**Figure 4F**), suggesting that transfections are independent stochastic events, though the rate of transfection likely varies over time. The mean transfection time was ∼140 min post transfection (mpt), with events scattered throughout the time we imaged. This time scale is consistent with the fast kinetics of mRNA transfection and translation (Leonhardt et al., 2014). Most cells reached their maximal luminescence rapidly post transfection, whereas a few cells grew luminescent over a longer period (**Figure 4G**). To quantify the rate of translation, we measured the difference between the time when luminescence maximizes (t_max_) and the time at which transfection occurs (t_0_) and found that on average cells reached their maximum luminescence within 60 min after transfection (**Figure 4H**).

Over time, we noticed that the luminescence in individual cells never plateaued but instead became dimmer after the peak over the course of a few hours regardless of the transfection time (**Figure 4E**, **Video S2**). This phenomenon is unexpected as even if mRNA, once delivered, is rapidly degraded within the cytoplasm, Nluc protein is reported to have a half-life of ∼8 days in cell lysates (Hall et al., 2012). In addition, Nluc is an ATP-independent luciferase, and therefore should not depend on cell metabolism to luminesce. We wondered whether the dimming was primarily due to substrate depletion, so we added additional substrate after the first imaging session when only a few dimly luminescent cells remained. While some cells indeed recovered partially, most cells remained dark (**Figure S9**), suggesting that substrate availability was not the limiting factor. Notably, we occasionally saw cells burst open, releasing Nluc into the media. This was evident from the single-cell luminescence showing a sudden spike and an immediate drop at the moment of rupture (**Figure 4I**). However, these events were rare during our imaging experiments, so cell death was not the major cause of luminescence decay either. These results implied that either planarian cells have special mechanisms that caused Nluc to be particularly unstable or sequestered, or Nluc was released, actively via secretion or passively through leaky membranes, though the exact mechanism requires further investigation. Together, these results suggest that it is feasible to image Nluc in live cells at high temporal resolution, which provides a powerful tool for quantifying gene expression kinetics in planarian cells.

### mRNA transfection can reveal post-transcriptional regulatory elements

The ability to transfect and express exogenous mRNA also opens new possibilities for the analysis of gene regulation in planarians, which have so far been limited to observing phenotypic and measuring genotypic changes under the influence of RNAi. For example, the synthesis of an mRNA and the abundance of its encoded protein can be decoupled by cis-acting elements in the 5’ and 3’ UTRs that affect translation or mRNA stability. Understanding which regions of a UTR sequence are responsible for such post-transcriptional regulation requires the ability to remove or add putative regulatory elements to a reporter gene and study how its expression is altered.

As a case study, we selected a transcript (dd_Smed_v6_62_0_1) which contains two long overlapping open reading frames (ORF1: 513 nt, ORF2: 483 nt) with ORF1 beginning 40 nt upstream of ORF2. To resolve which ORF (or both) is translated, we created two constructs with Nluc beginning at the start codon of each reading frame. Transfecting these reporters into bulk dissociated cells showed that the translation was specific to the first open reading frame (ORF1), as Nluc expression was barely detectable from the ORF2 (**Figure 5A**). Removing the start codon of ORF1 by replacing the 5’ UTR with a synthetic sequence (attP1) recovered the Nluc expression. Thus, translation of these transfected RNAs was consistently initiated from the upstream start codon.

**Figure 5:**
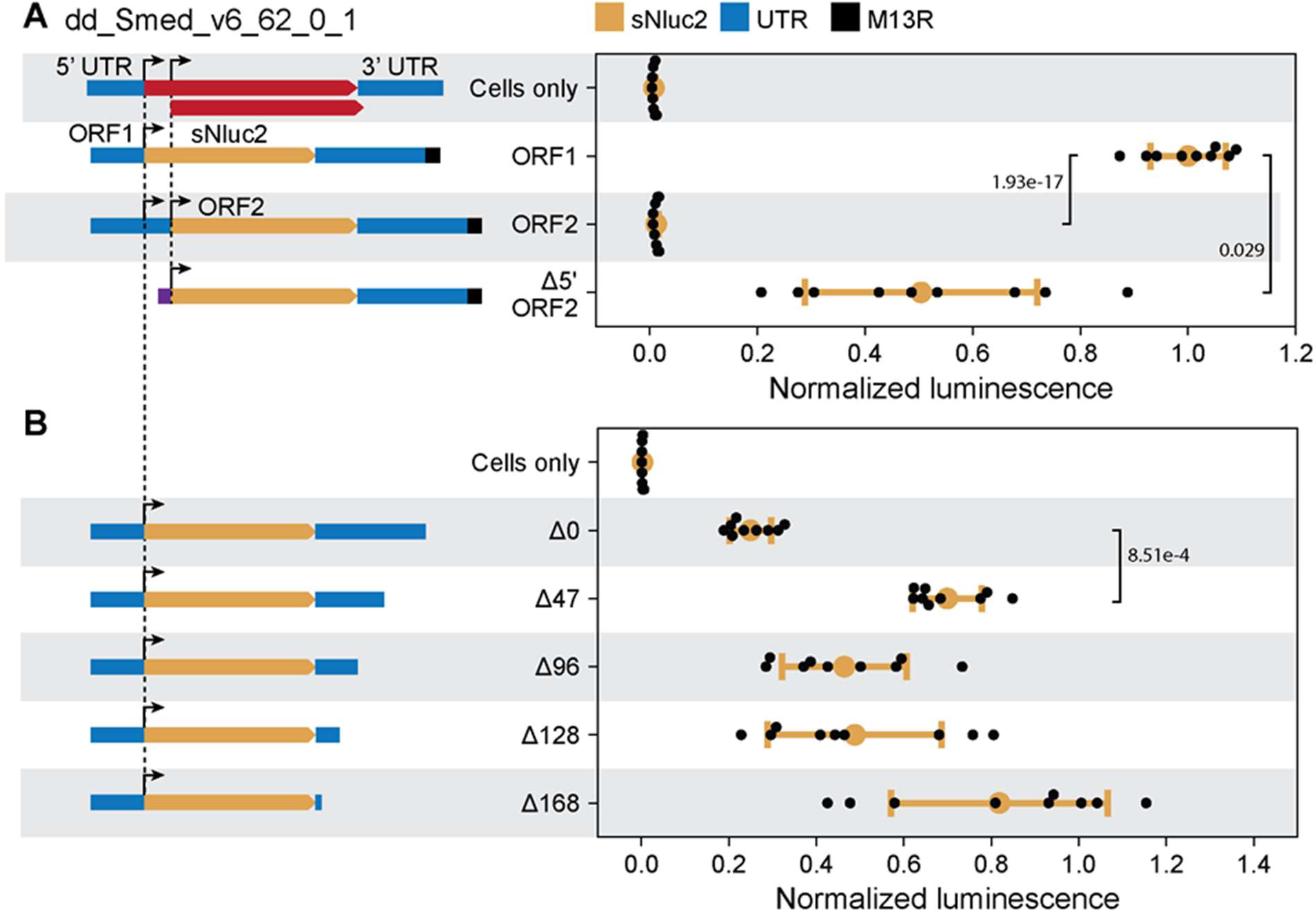
mRNA transfection enables studies of post-transcriptional regulation. **(A)** Luminescence is detected with Nluc coding sequence beginning at ORF1 but not at ORF2 of dd_Smed_v6_62_0_1. Replacing the full-length 5’ UTR of ORF2 with a synthetic UTR consisting of the attP1 sequence rescues the expression from ORF2. Note that there is a M13R synthetic sequence at the 3’ end for all three constructs. In schematics, the dashed lines mark the beginning of ORFs. Luminescence is measured at room temperature with NanoGlo-Live substrate. All transfections in this figure are performed with 200,000 cells from whole dissociated planarians, using 0.8 µL Viromer and 1 µg mRNA per well, and assayed 24 at hpt. **(B)** Expression from the ORF1 construct with the 3’ UTR successively truncated. Statistics. All data points presented are technical replicates (n = 3) from independent biological replicates (n = 3). All values reported are normalized to the expression of ORF1-sNluc2. Error bars: standard deviation. ANOVA: p = 1.83e-17 (**A**) p = 3.36e-7 (**B**). p-values for pairwise comparisons are calculated using two-sided Welch’s t-test and reported in the figure.

Next, we investigated the function of the 3’ UTR. For this, we successively truncated the 3’ UTR from the ORF1 Nluc construct. We found that removing the end of the 3’ UTR, nucleotides 141-188 (Δ47), increased the expression by 3-fold, suggesting the presence of a destabilizing or repressing element within this region; deleting additional sequences had minimal additional effects (**Figure 5B**). Surprisingly, adding a short synthetic sequence (M13R) to the 3’ end of the full-length UTR drove considerably stronger expression (**Figure 5A**), implying that the function of the putative repressive or destabilizing elements may be sensitive to sequence context. Overall, these results provide a platform for studying post-transcriptional regulation in planarian cells.

### Heterologous reporter expression in live animals

Although the ability to transfect planarian cells with mRNA *in vitro* represents a significant advance in the genetic manipulation of planarian cells, current planarian cell culture is limited by low cell viability, lack of cell proliferation, and the gradual loss of neoblast identity (Kai et al., 2019), thus limiting the range of questions that can be addressed in isolated cells. We therefore explored whether our transfection protocols might be sufficient for generating reporter expression in whole animals.

We injected RPL15-sNluc2 mRNA complexed with Viromer or Trans-IT into the parenchymal tissue along the tail midline, which reduces the risk of misinjections into the abundant gut branches (**Figure 6A**). We then measured bulk luminescence on individually dissociated worms at multiple time points post transfection. The luminescence background in tissue lysates from sham-injected animals was universally very low, mirroring what was observed *in vitro*. In contrast, mRNA injection with Viromer or Trans-IT led to luminescence signals up to 1000-fold above background (**Figure 6B-C**). For Viromer, the highest fraction of expressing animals was detected already at 2 hpt (30/30) and remained robust for 12 hpt (29/30) (**Figure 6B**). Contrary to our observations *in vitro*, Trans-IT transfections produced luminescence intensities stronger than those of Viromer transfections by an order of magnitude (**Figure 6C**), suggesting that for *in vivo* transfections, Trans-IT is more effective than Viromer.

**Figure 6:**
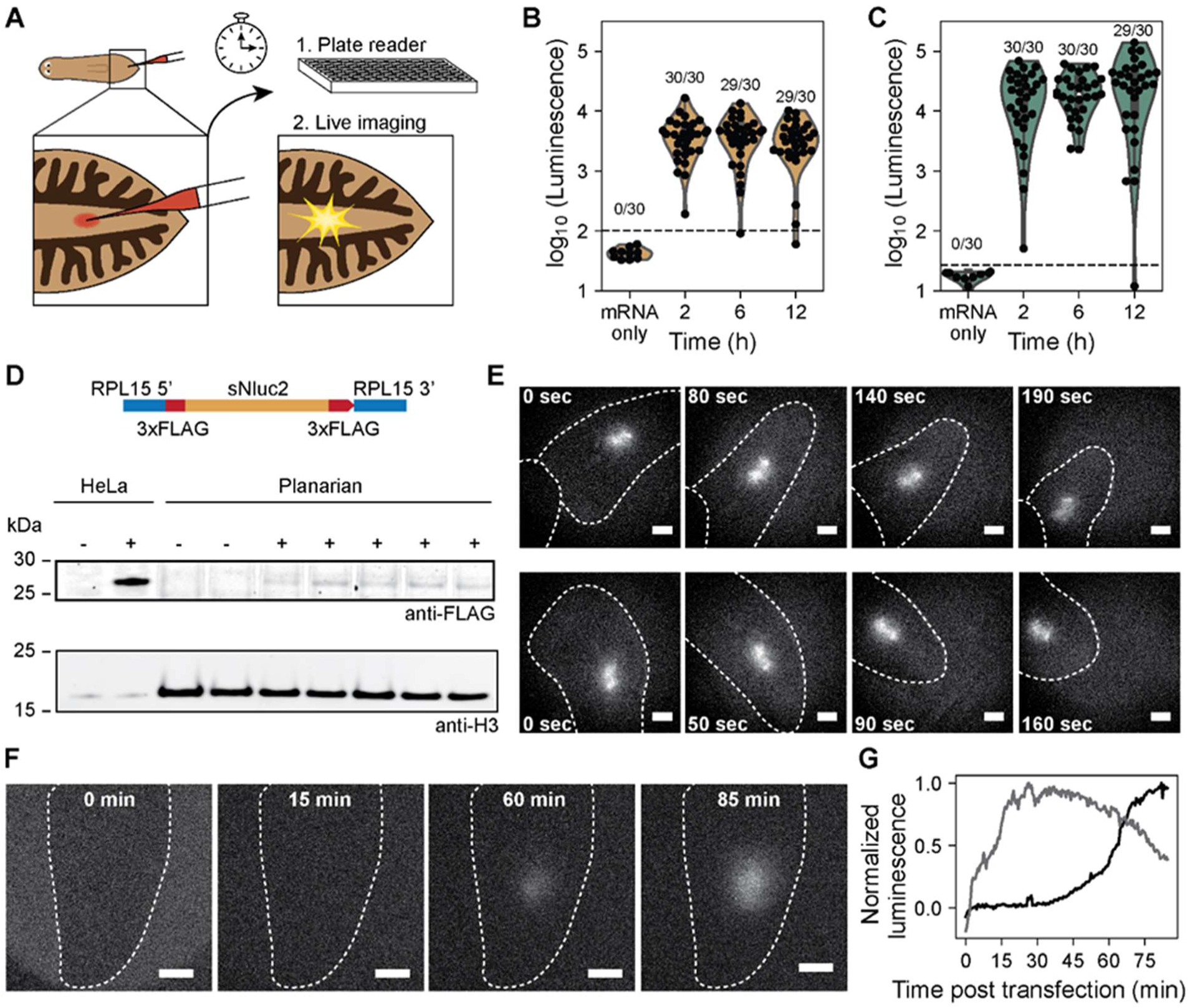
Nluc expression in live animals. **(A)** Schematics showing the workflow of *in vivo* transfection experiments. (**B-C**) Bulk luminescence is measured from dissociated tissues injected with mRNA complexed with Viromer mRNA (**B**) or Trans-IT mRNA (**C**). Dashed lines: 3 standard deviations above the background (based on the mRNA only condition) to discriminate negative and positive animals. Numbers of positive animals out of all animals injected are reported for each time point. Transfection complexes contain either 0.8 µL of Viromer, 1 µg of RPL15-sNluc2 mRNA, and Viromer Buffer up to a final reaction volume of 25 µL, or 2 µL of Trans-IT, 1 µL of Boost, 1.5 µg of RPL15-sNluc2 mRNA, up to a final volume of 25 µL in L-15. For each transfection reagent, 30 animals, from two independent experiments, are injected along the tail midline with 0.9 µL of transfection mix per time point. Transfected animals are dissociated, and luminescence is measured at room temperature using NanoGlo lysis reagent. **(D)** Western blot of Nluc protein from Trans-IT transfected planarian lysates. Animals are injected along the tail midline and amputated at 12 hpt to collect the tails. Three tails are pooled for each sample, from which protein is extracted. Hela cells are transfected with Trans-IT using the same condition. Top: schematic diagram of the 6xFLAG-sNluc2 construct. **(E)** Snapshots from a time-lapse video of two Trans-IT transfected animals imaged at 4 hpt. Images are collected at 1 frame per min using 10 s exposure time and gain of 500 on an Andor iXon DU-897 EMCCD with a 10× air objective (BoliOptics, N.A. = 0.25). Animals are decapitated to reduce motility. Dashed lines: animal body boundary. Scale bars, 200 µm. **(F)** Expression kinetics of *in vivo* transfection. Animals are transfected using Trans-IT and allowed to rest for 30 min. Undiluted FFz substrate is injected into the gut and the animal is embedded in anesthetic agarose solution (2% low melting point agarose, 0.1% chloretone, 1:5000 linalool) and imaged. Images show the luminescence intensity before (left) and during (right) expression. Dashed lines: animal body boundary. Scale bars, 200 µm. **(G)** Time traces of *in vivo* luminescence from two independently injected animals. Luminescence is quantified as the difference between the total intensities from two equal-sized regions centered at and above the injection site to subtract out ambient light intensity. The data is normalized by the maximum luminescence.

To confirm that luminescence came from cells, we dissociated transfected tails, and through imaging, we observed high densities of luminescent cells (**Figure S10**). Further, we succeeded in validating Nluc expression using Western blotting. Initial efforts using an anti-Nluc antibody failed due to poor binding characteristics; as a result, we transfected planarians with RPL15-sNluc2 flanked by two 3× FLAG affinity tags. Western blotting against anti-FLAG produced a clear band, providing an orthogonal validation of Nluc expression (**Figure 6D**). As a comparison, an equivalent mass of transfected HeLa cell lysate was also blotted, which produced a significantly brighter signal consistent with the higher expression observed in luminescence assays. Together, these results establish mRNA reporter delivery and expression in live planarians.

We next asked whether Nluc expression was sufficient for luminescence imaging *in vivo*. We injected at either the tail midline or behind the left eye and incubated the animals for 15 min in NanoGlo-Live substrate supplemented with 1% DMSO to aid in tissue penetration. The immobilized animals were then imaged on an LV200 (Olympus) microscope. This experiment revealed bright luminescence after both Viromer and Trans-IT injections, while no luminescence was detected in negative controls injected with mRNA alone (**Figure S11**). Although the spatial resolution of *in vivo* luminescence imaging did not allow for cellular resolution, the small size of the luminescent region was consistent with the transfection of a small cluster of cells around the injection site.

Luminescent reporters are particularly attractive for live imaging of planarians, as these animals are photophobic and become agitated by the intense excitation illuminations required for fluorescence experiments. Therefore, we tested time-lapse imaging of unrestrained animals and succeeded in tracking luminescent patches between frames (**Figure 6E**, **Video S3**). For these experiments, we used FFz as a low toxicity substrate and found that the animals can be recovered alive after a few hours of imaging. Encouraged by this result, we then imaged planarians right after injecting Trans-IT-mRNA complexes to determine expression kinetics *in vivo.* We further increased the luminescence signal by injecting the FFz substrate directly into the planarian gut to improve the bioavailability and to reduce the amount of FFz used per experiment. By continuously imaging for ∼1 hr, we observed similarly rapid expression kinetics consistent with our observations *in vitro* (**Figure 6F-G**, **Video S4**). These results represent the first direct measurement of gene expression kinetics in a live planarian, which establishes luminescence as a route for quantitative live imaging through thick, strongly autofluorescent tissues.

## Discussion

Transgene expression in planarian flatworms has been a persistent challenge in the field since the molecular biology revival of the system around two decades ago (Agata & Watanabe, 1999; Newmark & Sánchez Alvarado, 2002). Here, we accomplished heterologous protein expression in the planarian model species *S. mediterranea* by combining four experimental approaches: (1) nanostraw electro-delivery to establish an initial positive control, which enabled subsequent optimization of chemical delivery methods; (2) delivering mRNA instead of DNA to bypass the complexities of nuclear import, transcription, and splicing; (3) optimizing post-transcriptional factors to enhance expression; and (4) using a luminescent reporter to circumvent the strong autofluorescence which complicates the use of fluorescent reporters. By making these choices, we observed a clear signal from exogenously supplied Nluc mRNAs in planarian cells both *in vitro* and *in vivo*.

Exogenous reporter expression in our results is supported by multiple independent lines of evidence. First, we showed that planarians are minimally autoluminescent, and most experimental conditions tested reproducibly reached luminescence intensities as high as 3-4 orders of magnitude above the background, which makes the detection of false positives extremely unlikely. Second, we observed that luminescence showed a dose-dependence on Nluc mRNA amounts, which was abolished by the addition of a premature stop codon. Third, luminescence intensity was modulated by biologically relevant factors such as UTR sequences and codon usage bias. Fourth, imaging of transfected cell cultures confirmed a cytoplasmic origin of the luminescence signal, and time-lapse imaging showed rapid expression kinetics in single cells consistent with mRNA expression. Fifth, we succeeded in imaging luminescence in live injected animals, with the signal consistently restricted to the injection site. Sixth, we were able to detect Nluc protein via Western blotting from tissue lysates of transfected animals. Finally, the results presented in this manuscript were gathered in two different laboratories, which highlights the robustness and general applicability of the technique. Collectively, our results represent the first unambiguous demonstration of exogenous mRNA expression in planarians.

In terms of direct utility, our current reporter assay represents a significant expansion of the planarian toolkit. We demonstrated that by delivering Nluc mRNA, we were able to identify successful transfection methods and subsequently optimize them. New gene delivery strategies are being developed at a rapid pace, including dendrimers, nanoparticles, virus-like particles (VLPs), and cell-penetrating peptides (Lostalé-Seijo & Montenegro, 2018). Our methodology can be continuously applied to screen this ever-expanding repertoire of transfection techniques to achieve more efficient delivery, lower cytotoxicity, and higher expression. Similarly, this approach can be used to assess other mRNA-based expression systems such as self-replicating RNA replicons including alphavirus (Beal et al., 2015) and nodamuravirus (Taning et al., 2018), which may help to overcome mRNA’s transient expression and increase overall luminescence intensity. In even broader contexts, our method also allows testing additional reporters and transfecting other species, with initial success reported in **Figure S12** and **Figure S13**, respectively.

Importantly, the use of FFz, a novel water-soluble furimazine derivative, allowed us to monitor the kinetics of transfection and expression *in vitro* and *in vivo* by time-lapse luminescence imaging. This technique could be used not only to assess novel transfection reagents for favorable kinetics, but also to monitor the dynamics of biologically relevant processes occurring on the timescale of minutes to hours. Luminescent reporters have two clear advantages over fluorescent reporters for live imaging applications. First, planarians are photophobic and escape quickly from the bright excitation lights used in fluorescence imaging. Second, planarians are highly autofluorescent, which makes interpreting weak fluorescence signal difficult if not impossible. With luminescence, we detected Nluc expression in unrestrained animals and captured initial expression kinetics on live animals immediately after transfection. We anticipate that the ability to sensitively detect Nluc expression *in vivo* will be valuable for screening other expression systems, such as DNA-based transgenesis, especially since animals may be recovered alive after luminescence imaging.

Beyond the technical applications, we demonstrated that Nluc mRNA transfections can be used to study post-transcriptional regulation, a biological application that we anticipate will be of broad interest. In particular, we quantitatively measured the consequences of varying UTR sequences. This approach provides a route to identify novel cis-regulatory sequences, which would be impossible otherwise. Such sequences may serve as tools for further controlling gene expression through a broader dynamic range. Besides UTRs, our method allows for exploring other post-transcriptional regulatory mechanisms in planarians, such as incorporating trans-spliced 5’ leader sequences (Zayas et al., 2005; Rossi et al., 2014), adding target sites of known small RNAs (Kim et al., 2019), including internal ribosomal entry sites (IRES), or manipulating secondary structures of mRNA (Leppek et al., 2022). Our study sets the stage for the systematic characterization and analysis of these factors to better predict how sequence informs expression. In addition to regulatory sequences, the individual nucleotides in mRNA can be modified to increase RNA expression and/or stability (Andries et al., 2015; Svitkin et al., 2017). As a first attempt, we synthesized mRNA incorporating the canonical uridine versus mRNA containing the base analog 1-methyl pseudouridine (m1Ψ) and found that both RNAs were capable of expression in planarians. While we did not observe any significant differences in expression levels (**Figure S14**), our method provides a route to explore the effects of other modified nucleotides on mRNA expression in planarian cells.

Achieving planarian transgenesis will require solving numerous challenges, including gene delivery, expression, and detection. Our method addresses them by utilizing several transfection reagents to deliver a Nluc reporter construct, the expression of which we can unambiguously detect and quantitatively measure. Collectively, these narrow the enormous parameter space that makes the *de novo* establishment of transgenesis in phylogenetically distant model systems so challenging. As such, our results transform the quest for planarian transgenesis from a trial-and-error process into a constrained parameter optimization process, which we anticipate proceeding rapidly with our community. More broadly, we predict that our strategies may also prove useful for manipulating other asexually reproducing animals which lack a germline and cannot otherwise be transformed through embryonic manipulations.

### Limitations of Study

Compared to the current state of the art in established genetic model systems, the absolute level of reporter expression is relatively low in our study. Although this limits the use of fluorescent reporters (especially given the strong autofluorescence of planarian tissues), we found that Nluc enabled the development of a much higher signal-to-noise reporter, allowing sensitive and quantitative detection of gene expression, especially when used with improved water-soluble substrates. In addition, the chemical transfection reagents were screened and optimized on total cells, which may limit their efficacy and specificity on stem cells (neoblasts). While we detected Nluc expression in neoblast-enriched cell populations, we could not rule out that the signal may originate from a small fraction of co-purified differentiated cells. We also performed *in situ* hybridization against the neoblast marker *piwi-1* on cells dissociated from transfected animals but failed to detect convincing cases of Nluc/*piwi-1* co-localization. Hence it is possible that transfections of neoblasts are rare, or that upon transfection, neoblasts might lose *piwi-1* expression, which is known to be sensitive to the cell cycle (Raz et al., 2021). These results suggest that efforts need to be made either to identify transfection methods that are more efficient at transfecting neoblasts specifically, or to better understand how transfection perturbs neoblast identity. Finally, of the two most effective transfection reagents, one (Viromer mRNA) is no longer commercially available. Since Trans-IT and Viromer are both cationic polymers, other reagents with similar chemistries may provide substitutes for *in vitro* transfections. For *in vivo* transfections, Trans-IT is most effective, remains commercially available, and thereby represents the most promising route going forward towards planarian transgenesis.

## Acknowledgements

We thank K. Lei and C. Kuhn for stimulating discussion, E. Liu, D. Burghardt, M. Khariton, B. Rosental, Y. Xue, Y. Lim, and T. Boothe for technical assistance. R.N.H. is supported by an NSF Graduate Research Fellowship. U.W. is supported by an EMBO Long-Term Fellowship. B.W. is Beckman Young Investigator. U.W., L.D., and J.C.R. are supported by the Max Planck Society. A.Z.F is supported by a NIH grant (R35GM130366). This work was initiated through a Stanford Bio-X Interdisciplinary Initiative seed grant (IIP9-64) to B.W., A.Z.F., and N.A.M., and later supported by an NSF EDGE grant (IOS-1923534) to B.W., A.S.A., and N.A.M., and a Volkswagen Foundation grant (Az.-99498) to J.C.R. and B.W.

## Author Contributions

Conceptualization, R.N.H., U.W., A.F., J.C.R., and B.W.; Methodology, R.N.H., U.W., L.D., S.L.O., and H.L.; Investigation, R.N.H., U.W., L.D., and C.C.; Writing – Original Draft, R.N.H., J.C.R, and B.W.; Writing – Review & Editing, R.N.H., U.W., S.L.O., A.S.A., A.Z.F., J.C.R., and B.W., with input from all other authors; Supervision, N.A.M., A.Z.F., J.C.R, and B.W.; Funding acquisition: A.S.A., N.A.M., A.Z.F., J.C.R, and B.W.

## Resource Availability

### Lead Contact

Further information and requests for resources and reagents should be directed towards the lead contact, Bo Wang.

### Materials Availability

Plasmids used in this study are available freely upon request and have been deposited to Addgene (#186753-#186774, and #187221). Though no longer commercially available, Viromer mRNA is available upon request while our limited supplies last. Nanostraw electro-delivery is presently being commercialized by NAVAN Technologies, who can be reached for inquiry. All species and strains of planarians used in this study are available upon request.

### Data and Code Availability

- riginal Western blotting images, raw plate reader values, and microscopy data have been deposited on Figshare.
- data and code have been deposited on Figshare, and our GitHub accounts. DOIs are listed in the key resources table.
- additional information required to reanalyze the data reported in this paper is available from the lead contact upon request.

### Experimental Models and Subject Details

#### Planarian Models

Asexual *S. mediterranea* strain CIW4 were reared in the dark at 20 °C and maintained in 0.5 g/L Instant Ocean (Carolina Biological Supply, Cat#671442) supplemented with 0.1 g/L sodium bicarbonate and were fed a diet of macerated calf liver once per week. *S. polychroa* strains were reared in the dark at 20 °C and maintained in 0.75× Montjuic Salts (1.2 mM NaCl, 0.75 mM CaCl_2_, 0.75 mM MgSO4, 0.075 mM MgCl_2_, 0.075 mM KCl) supplemented with 0.1 g/L sodium bicarbonate and were fed a diet of macerated calf liver once per week.

## Method Details

### Planarian cell dissociation

Planarian cells were prepared by finely mincing 10-15 asexual *S. mediterranea* (5-7 mm in length) with a razor blade and suspending the tissue in CMF (Ca/Mg-Free media: 480 mg/L NaH_2_PO_4_, 960 mg/L NaCl, 1.44 g/L KCl, 960 mg/L NaHCO_3_, 3.57 g/L HEPES, 0.24 g/L D-glucose, 1 g/L BSA, pH 7.4 in MilliQ H_2_O). The tissue was rocked for 5 min, followed by gentle pipetting for 10 min for 3 times, or until the tissue was visibly homogenized. The cells were centrifuged at 250 g for 4 min, and the supernatant was removed and replaced with 1.5 mL of fresh CMF. The cell suspension was then serially filtered through 100, 70, 40, and 35-μm mesh strainers. The filtered cell suspension was centrifuged and transferred to Iso-L15 (1:1 Leibovitz’s L-15 to MilliQ H_2_O, 1× MEM nonessential amino acids, 1× antibiotic-antimycotic, 1× MEM vitamin solution, 1 mM Sodium Pyruvate, 2.5 g/L HEPES, 5% FBS, buffer to pH 7.8).

### FACS

To isolate a neoblast-enriched X1FS population, an aliquot of sacrificial cells was stained with Hoechst 33342 (10 µg/mL, ThermoFisher, Cat#H3570) in CMF for 15 min, filtered, and sorted on a Sony SH800 with either the 100 or 130 µm sorting chip (Sony, Cat#LE-C3210, Cat#LE-C3213). Cells were gated for size using forward and side-scatter signals, then neoblasts were identified by gating, in linear scale, on cells with high Hoechst blue (Excitation: 405 nm, Emission 450/50 nm) and low Hoechst red (Excitation: 405 nm, Emission 600/60 nm) signal (Hayashi et al., 2006). Following the identification of the neoblast population using Hoechst fluorescence, unstained planarian cells were loaded and sorted into Iso-L15 medium using the X1FS gate overlaid on the forward and side-scatter. After sorting, the cells were centrifuged at 250 g for 5 min and resuspended in fresh Iso-L15.

### Nanostraw electro-delivery

Nanostraws were acquired from NAVAN Technologies. Each nanostraw cartridge was loaded with 200,000 X1FS cells in 300 µL of Iso-L15 media and centrifuged at 300 g for 10 min to ensure close contact between straws and cells. For each cartridge, 3 µg of *in vitro* synthesized mRNA was diluted in PBS to a total volume of 35 µL and placed on the titanium anode, and a cartridge was carefully lowered onto the mRNA solution. The titanium cathode was placed atop, and the electro-delivery assembly was subjected to a 35 V, 200 µs, 40 Hz square wave pulse 3 times for 45 s each, with a 1 min rest in between pulses. Transfected cells were incubated at 20 °C in the dark for 24 hr before being transferred from the nanostraw cartridge to an opaque white 96 well plate (Greiner, Cat#655075) for assaying luminescence.

### Chemical transfection

For *in vitro* experiments, dissociated planarian cells were suspended at a concentration of 0.88×10^6^ cells/mL in Iso-L15 medium supplemented with 10 µg/mL ciprofloxacin (Sigma-Aldrich, Cat#PHR1044). 225 µL of cell suspension was added to each well of white opaque (for plate-reader assays) or glass bottom (for luminescence imaging) 96-well plates for a total of approximately 200,000 cells per well. For the initial screen, each reagent was prepared as specified in **Table S2**, and for all subsequent experiments, we utilized the optimal reagent ratios identified in Figures 2C-D. After adding transfection complexes to each well, the cells were incubated at 20 °C in the dark for 4-72 hr before assaying luminescence. Biological replicates were performed by transfecting independently synthesized batches of mRNA, dissociated planarian cells, and assembled transfection complexes. Within each biological replicate, transfection mixes were split evenly across 3 parallel transfections of cells isolated from the same dissociation.

For live animal injections, transfection mixes, prepared in the same manner as *in vitro* reactions, were loaded into needles pulled from glass capillaries (WPI, Cat#1B00F-3) on a Sutter P90 needle puller with the following settings: pressure = 500, heat = 758, pull = 50, velocity = 70, time = 200. Needles were loaded on a FemtoJet injection system (Eppendorf) or Sutter XenoWorks injection system, and the needle tip was opened with forceps. Animals were placed ventral-side up on moist filter paper placed on a cooled block. Animals were injected along the tail midline with 900 nL transfection mix or until a bolus of injected fluid was visible and ceased expanding. For ocular injections, animals were placed ventral-side down and injected immediately posterior to the left eye cup. Animals were left to rest in the dark at 20 °C until assaying 4-24 hpt.

### Luciferase plate assay

For dissociated cells, luminescence was measured using the NanoGlo-Live Cell Assay (Promega, Cat# N2011). NanoGlo-Live substrate was added to NanoGlo buffer at a ratio of 1:20, and 25 µL of reagent was added to 250 µL of transfected cells. NanoGlo-Live reagent was also added to negative control conditions before assaying to ensure a consistent luminescence baseline.

For live planarians, each injected animal was individually dissociated by finely mincing with a razor blade and suspending the tissue in 250 µL Iso-L15 medium just prior to assaying. The resuspended tissue was transferred to an opaque white 96-well plate. Nluc expression was measured using the NanoGlo Luciferase Assay System (Promega, Cat# N1110). Substrate was added to the NanoGlo lysis buffer at a ratio of 1:50 and 100 µL of reagent was added to the cells. Cells were lysed by pipetting up and down 10 times or until the tissue was visibly homogenized.

Gaussia luciferase (Gluc) was assayed using the Pierce™ Gaussia Luciferase Glow Assay Kit (ThermoFisher, Cat#16160) according to the manufacturer’s instructions. To measure luminescence from supernatant and cellular pellets, 250 µL of supernatant was carefully transferred to a fresh 96-well plate, then the remaining cells were resuspended in 250 uL of CMF. The resuspended cells were then transferred to a fresh 1.5 mL Eppendorf tube and were centrifuged at 250 g for 4 min. The supernatant was discarded, and the cells were then resuspended in 250 µL of Iso-L15 and transferred to an opaque white 96-well plate.

Luminescence was measured on a plate reader (BioTek Synergy™ HTX for collecting data presented in Figures 1, 3, 5, S5, S8, S12, S14; BioTek Synergy™ Neo2 for Figures 2C-D, 6B-C, S3, S13; and EnVision Microplate Reader for Figure 2B). Integration time was set at 1 s and the digital gain was kept consistent on each instrument for all experiments. All assays were performed at room temperature and luminescence was measured quickly after substrate was added to all wells.

### Luminescence imaging setup

Luminescence imaging was performed on an LV200 Bioluminescence Imaging System (Olympus) equipped with a 20× air objective (Olympus: UPLXAPO20X, N.A. = 0.8), a 60× oil immersion objective (Olympus: UPLXCAPO60XO, N.A. = 1.42), and a liquid-cooled Hamamatsu C9100-24B EMCCD camera (1024×1024 pixels). All images taken with the LV200 were acquired with an exposure of 60 s and a gain of 300 unless otherwise specified.

Images were also acquired with a custom-built luminescence microscope, modified from Kim et al., 2017, which utilizes an Andor iXon DU-897 EMCCD camera (512×512 pixels), a HIKROBOT MVL-HF5024M-10MP 50mm tube lens, and either a 10× air (BoliOptics: BM13013331, N.A. = 0.25), a 20× air (BoliOptics: BM03023431, N.A. = 0.4), or a 100× oil (Carl Zeiss: 1084-514, N.A. = 1.45) objective. All images taken with the custom luminescence microscope were acquired with an exposure time of 30 s and gain of 500 unless otherwise specified. Micromanager 1.4 was used to operate the microscope.

### Luminescence imaging *in vitro*

Nluc expression was detected using the NanoGlo-Live Cell Assay. A 35 mm glass-bottom dish (WPI, Cat#FD3510-100) or a glass bottom 96-well plate (Cellvis, Cat#P96-1-N) were coated with 0.5 mg/mL Concanavalin A (Sigma-Aldrich, Cat# L7647) for 2 hr, washed, and air dried. Transfected cells were transferred to coated dishes in a total of 250 µL of Iso-L15 and allowed to adhere to the glass surface for 1 hr. NanoGlo-Live substrate was added to NanoGlo buffer at a ratio of 1:20, and 25 µL of reagent was added to 250 µL of transfected cells. Cells were imaged at room temperature with either an LV200 Bioluminescence Imaging System (Olympus) with a 60× oil immersion objective (Olympus), or the custom-built luminescence microscope equipped with a 100× oil immersion objective (Carl Zeiss).

Quantification of transfection efficiency and viability were measured by staining transfected cells with Calcein AM (1 µM final concentration), propidium iodide (PI) (1 µg/µL final concentration), and Hoechst 33342 (1 µg/µL final concentration). The cells were imaged on an LV200 with a 20× air objective (Olympus), and individual cells were classified using scanR Analysis 3.2.0 (Olympus). Transfected cells were classified as being positive for Calcein, Hoechst, and Nluc. Viability was quantified by classifying live cells as positive for both Calcein and Hoechst. Total cells were quantified by Hoechst positive nuclei.

### Luminescence imaging *in vivo*

For *in vivo* imaging, animals were incubated in 100 µL of 1× Instant Ocean supplemented with 1% DMSO and 1:20 NanoGlo-Live substrate for 15 min at room temperature. The animals were then placed on glass bottom dishes cooled on ice. Excess water was wicked away, and the animals were embedded in 1.5-2% low-melt agarose gel (ThermoFisher, Cat#16520050) supplemented with 1:5000 linalool (Sigma-Aldrich, Cat#L2602) and 1:20 NanoGlo-Live substrate. A coverslip was placed over the agarose to prevent lensing. The animals were imaged using an LV200 with a 20× air objective (Olympus) at an exposure of 60 s and gain of 300.

### Time-lapse imaging *in vitro*

Bulk dissociated planarian cells were transfected with 0.8 µL of Viromer and 1 µg RPL15-sNluc2 mRNA. Cells were either imaged immediately or allowed to incubate for 24 hr at 20 °C. Substrate was prepared by diluting NanoGlo In-Vivo substrate (FFz) (Promega, Cat#CS320501) 1:50 in Iso-L15, then 25 µL of this solution was added to the 250 µL of transfected cell media. Cells were imaged at room temperature on a custom-built luminescence microscope with a 20× air objective (BoliOptics, N.A. = 0.4) at 1 frame per min with an exposure of 30 s and gain of 500. Individual cells were automatically segmented using Python 3.8 and scikit-image (see Code and Data Availability).

### Time-lapse imaging *in vivo*

To image unrestrained animals, they were allowed to incubate at 20 °C for 4 hr post injection, then amputated immediately anterior to the injection site. The animals were placed in a solution of 1× Instant Ocean containing 1:20 NanoGlo In-Vivo substrate (FFz) and imaged at a rate of 6 frames per min with an exposure of 10 s and gain of 500.

For measuring expression kinetics *in vivo*, animals were allowed to rest for 15 min post injection and re-mounted ventral-side up on moist filter paper over a cool block. Undiluted FFz was injected into the gut until at least one posterior gut branch had visibly filled with yellow substrate, then the animal was placed ventral-side down on a cooled glass-bottom dish. Excess water was wicked away before immobilization. Anesthetic agarose was prepared by mixing 2% agarose in 1% chloretone (Sigma-Aldrich Cat# 112054) dissolved in 1× Instant Ocean supplemented with 1:5000 linalool. The agarose solution was cooled to 37 °C and allowed to cool further just before gelling began, and drops were added on and around the animal to immobilize it. Animals were imaged at room temperature on a custom-built luminescence microscope with a 10× air objective (BoliOptics, N.A. = 0.25) at 2 frames per min with an exposure of 30 s and gain of 500.

### Cloning

To generate a plasmid for *in vitro* transcription and harvesting the UTR sequences, we first amplified the backbone of pDONOR221 (ThermoFisher, Cat#12536017) using primers BW-NH-104-105 (all primer sequences are provided in **Table S4**), as well as the LacZ cassette from pUC19 (Addgene #50005) with a T7 promoter sequence followed by a BsaI restriction site for subsequent cloning steps, all flanked between BbsI restriction sites and M13 forward and M13 reverse primer sites (for *in vitro* transcription template production) using primers BW-NH-106-107. The amplified backbone was digested with BsaI-HFv2 (NEB, Cat#R3733S) and the LacZ insert was digested with BbsI-HF (NEB, Cat#R3539S). The digested backbones were purified using the Zymo Clean and Concentrate kit (Zymo, Cat#D4004). The purified fragments were ligated together using T4 DNA Ligase (NEB, Cat#M0202S) to create pNHT7 (**Figure S6A**).

We cloned the gene of interest (GOI) from a pool of planarian cDNA using primers (BW-NH-108-125) containing BsaI restriction sites to produce overhangs compatible with pNHT7. The amplicons were purified using the Zymo Clean and Concentrate kit and then inserted into pNHT7 via a golden gate reaction containing 40 ng of backbone and 20 ng for each insert to be cloned in a 20 µL reaction volume containing 2 µL T4 Ligase Buffer, 1 µL T4 DNA Ligase, and 1 µL BsaI-HFv2 to produce pNHT7::GOI (**Figure S6B**).

Finally, to insert a reporter between the 5’ and 3’ UTRs, we amplified pNHT7::GOI with outward facing primers (BW-NH-128-139) which bind to the end and beginning of the 5’ and 3’ UTRs respectively containing the BsaI restriction sites. The reporter was then amplified to append compatible BsaI restriction sites using primers BW-NH-174-185 and inserted between the two UTR sequences via a golden gate reaction. The resulting plasmids were amplified with M13 forward and M13 reverse primers to produce linear template for *in vitro* transcription reactions (**Figure S6C**).

### *In vitro* transcription

Linearized templates for *in vitro* transcription were amplified using Phusion Polymerase (ThermoFisher, Cat#F531L) in two parallel 50 µL format reactions containing 10 µM M13F/R primers (**Table S4**) and 25 ng of template DNA. The two reactions were pooled and purified using the Zymo Clean and Concentrator kit, and the templates were eluted in 8 µL of RNase-free water. For the ‘UTR hacking’ experiments (Figure 5), PCRs were performed as described but replacing the M13R primer for an oligo which primes directly to the 3’ UTR (**Table S4**). Expected concentrations should range from 200-300 ng/µL.

*In vitro* transcription (IVT) was performed using the T7 mScript™ Standard mRNA Production System (CELLSCRIPT, Cat#C-MSC100625) according to the manufacturer’s protocol, opting for a 1.5 hr incubation during T7 transcription, a 2 hr incubation for 5’ capping, and a 1 hr incubation for poly-A tailing, all performed at 37 °C. RNA purification was performed by adding 600 µL of ethanol and 50 µL of 10 M ammonium acetate to 200 µL of IVT reaction, and allowing to precipitate overnight at -20 °C. The precipitated mRNA is then centrifuged at 21,000 g for 30 min at 4 °C. The pellet was then rinsed twice with 70% ethanol and allowed to air dry before being resuspended in 60 µL of nuclease-free water. A standard 60 µL reaction typically yields 60 µg of mRNA. For expected results, see **Figure S7**. For mRNA containing m1Ψ, the rNTP mix provided in the CELLSCRIPT kit was substituted for a mixture of rNTPs containing 10 mM rGTP, 10 mM rCTP, 10 mM rATP, and 10 mM m1Ψ (TriLink, Cat#N-1019-1). The IVT reaction can be scaled down by a half, though precipitation was done with the full volumes described here.

### Preparation of protein lysates

At 12 hpt, animals were washed twice with 1× Instant Ocean and decapitated to enrich for the injected regions. Three tails were pooled for each sample, transferred to a 1.5 mL tube and 80 µL Urea lysis buffer (9 M Urea, 100 mM NaH_2_PO_4_, 10 mM Tris-Base, 2 % w/v SDS, 130 mM DTT, 1mM MgCl_2_) was added. Animals were immediately lysed with a motorized pestle, followed by incubation at room temperature for 20 min to fully denature proteins. Cellular debris was then pelleted by centrifugation at 21,000 g for 15 min and supernatant was moved to a fresh 1.5 mL tube. Small aliquots from the protein lysates were diluted 1:10 in MilliQ H_2_O to quantify protein concentration based on 280 nm absorbance using a spectrophotometer. 20 µL of 5× LDS buffer (530 mM Tris HCl, 700 mM Tris-Base, 10 % w/v LDS, 50 % w/v glycerol, 2.55 mM EDTA, 500 mM DTT, 0.11 mM SERVA Blue G250, 0.875 mM Phenol Red, pH 8.5) was then added to each sample.

### SDS-PAGE and Immunoblotting

SDS-PAGE and Western blotting were performed using a XCell SureLock Mini-Cell and XCell II system (Invitrogen, Cat#EI0002). 40 µg total protein from planarian lysate or 5 µg total protein from HeLa cell lysate were loaded per well onto a NuPAGE 4-20 % BisTris gel (ThermoFisher, Cat#NP0321BOX) and run in 1× MOPS buffer (ThermoFisher, Cat#NP0001) at 125 V for 110 min. Proteins were blotted onto a 0.45 µm nitrocellulose membrane (Merck, Amersham Protran, Cat#GE10600002) for 2 hr at 4 °C and 30 V. Membranes were washed with PBSTw (PBS supplemented with 0.1 % Tween 20) twice followed by blocking for 1 hr in PBS with 5 % (w/v) soy protein isolate (Powerstar Food, Cat#psf-1131). Membranes were then incubated overnight at 4 °C on a horizontal shaker with primary antibody in PBSTw supplemented with 0.5 % (w/v) soy protein isolate. Membranes were then washed 4 times with PBSTw over 1 hr and incubated with secondary antibody in PBSTw with 5 % (w/v) soy protein isolate for 2 hr on a horizontal shaker. Membranes were again washed 4 times with PBSTw over 1 hr, rinsed with PBS twice and dried for 1 hr. Images were acquired on an Amersham Typhoon imaging system (Cytiva).

Mouse anti-FLAG-M2 (Merck, Cat#F3165; dilution 1:5000) and Rabbit anti-Histone3 (abcam Cat#ab1791; dilution 1:30000) were used as primary antibodies. Goat anti-mouse-CF770 (Biotium, Cat#20077; dilution 1:10000) and goat anti-rabbit-CF680 (Biotium, Cat#20067; dilution 1:10000) were used as secondary antibodies.

## Quantification and Statistical Analysis

All statistical tests were performed using Python 3.8 and the SciPy 1.8.0 stats package in Jupyter notebooks which have been deposited to Figshare (see Data and Code Availability). Images were processed in ImageJ 1.53k, and single-cell segmentation and quantification was performed in scanR Analysis 3.2.0 (Olympus).

## Supplemental Information

### Supplemental Figures

**Figure S1:**
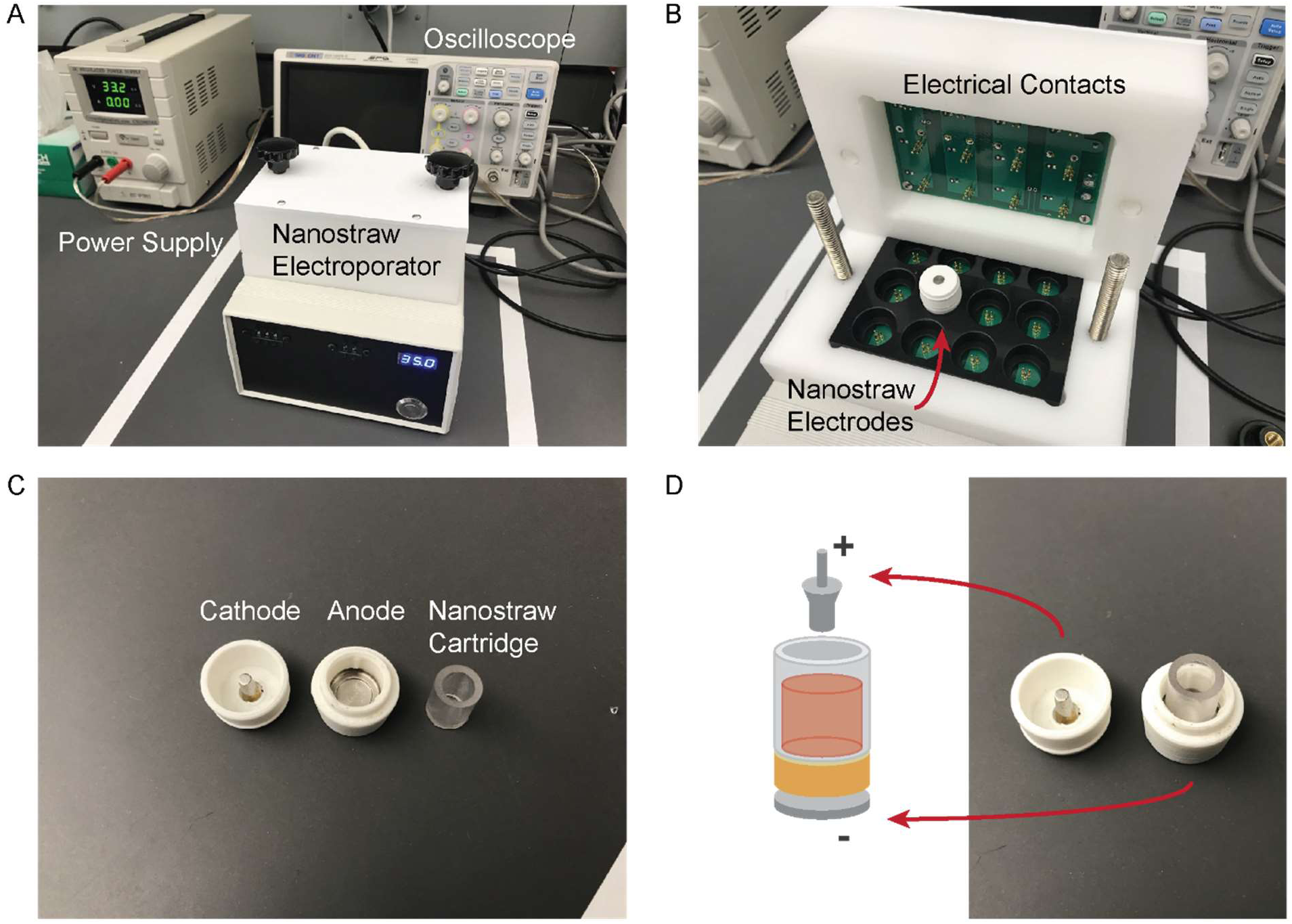
Nanostraw electro-delivery device, Related to Figure 1. **(A)** A photograph of the nanostraw device. In frame is the power supply (left), the oscilloscope, which is used to monitor the square wave pulses (top right), and the nanostraw electroporator device (center bottom) showing the dials to set the pulse width, duration, and voltage. **(B)** An interior view of the nanostraw device, showing the electrical contacts which interface with the nanostraw electro-delivery cartridge and carry the electrical pulses to the titanium electrodes on the top and bottom of the cartridge. **(C)** Individual components of the nanostraw cartridge, showing the cathode, which dips into the cell culture medium from above, the anode, which sits just below the buffer and mRNA to be delivered, and the nanostraw membrane, which is inserted atop the buffer and anode. **(D)** A demonstration of a nanostraw membrane placed within the cartridge.

**Figure S2:**
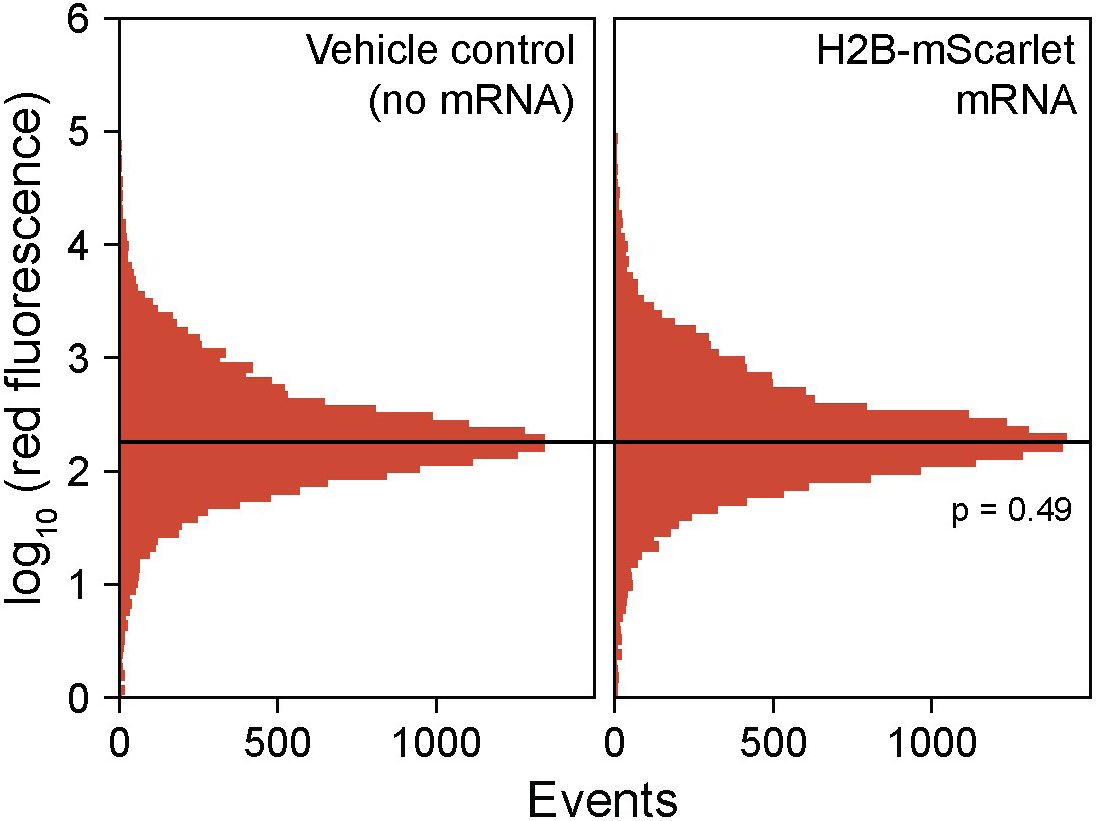
Autofluorescence in planarian cells, related to Figure 1. Histogram of red fluorescence (Excitation: 405 nm, Emission: 600/50 nm) of cells after nanostraw delivery measured by flow cytometry. Both samples show nearly identical distributions. Statistical insignificance (p = 0.49, two-sided Welch’s t-test) is established by bootstrapping, which calculates the p-value for 10,000 random subsamples of 100 cells between both distributions.

**Figure S3:**
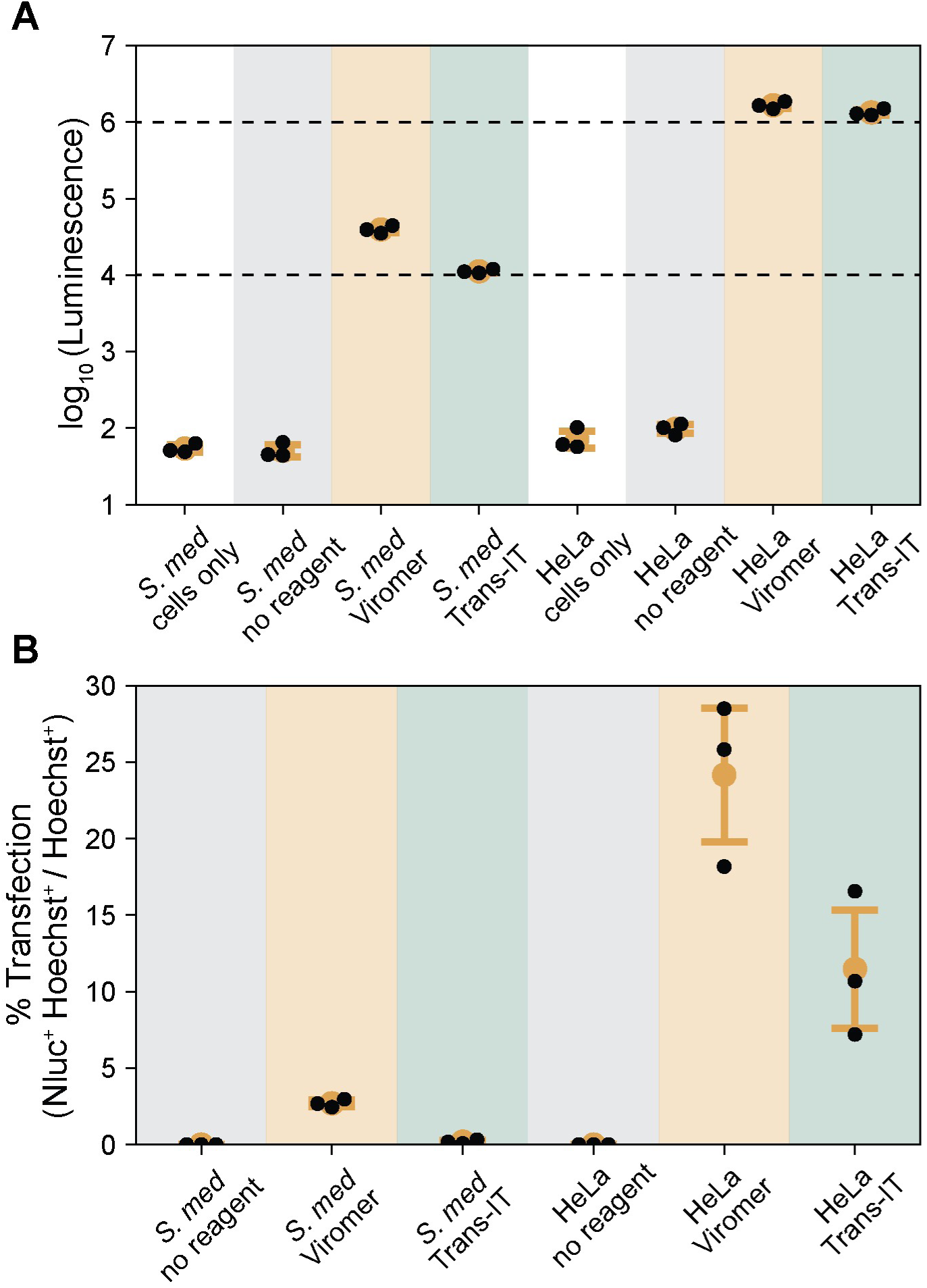
Comparisons of Nluc expression in planarian and HeLa cells, related to Figure 2. **(A)** Bulk luminescence of RPL15-sNluc2 transfected planarian and HeLa cells. All transfection are performed with 200,000 cells using optimal conditions identified in Figure 2C-D for planarian cells and **Table S2** for HeLa cells, and assayed 24 hpt at room temperature using NanoGlo-Live substrate. Dashed lines highlight 10^4^ and 10^6^ RLU. **(B)** Percent transfection efficiencies (Nluc^+^Hoechst^+^/Hoechst^+^) of transfected planarian and HeLa cells. Statistics. All data points presented are technical replicates using the same batch of mRNA. Error bars: standard deviation. ANOVA p = 1.35e-14 (**A**); p = 1.21e-6 (**B**).

**Figure S4:**
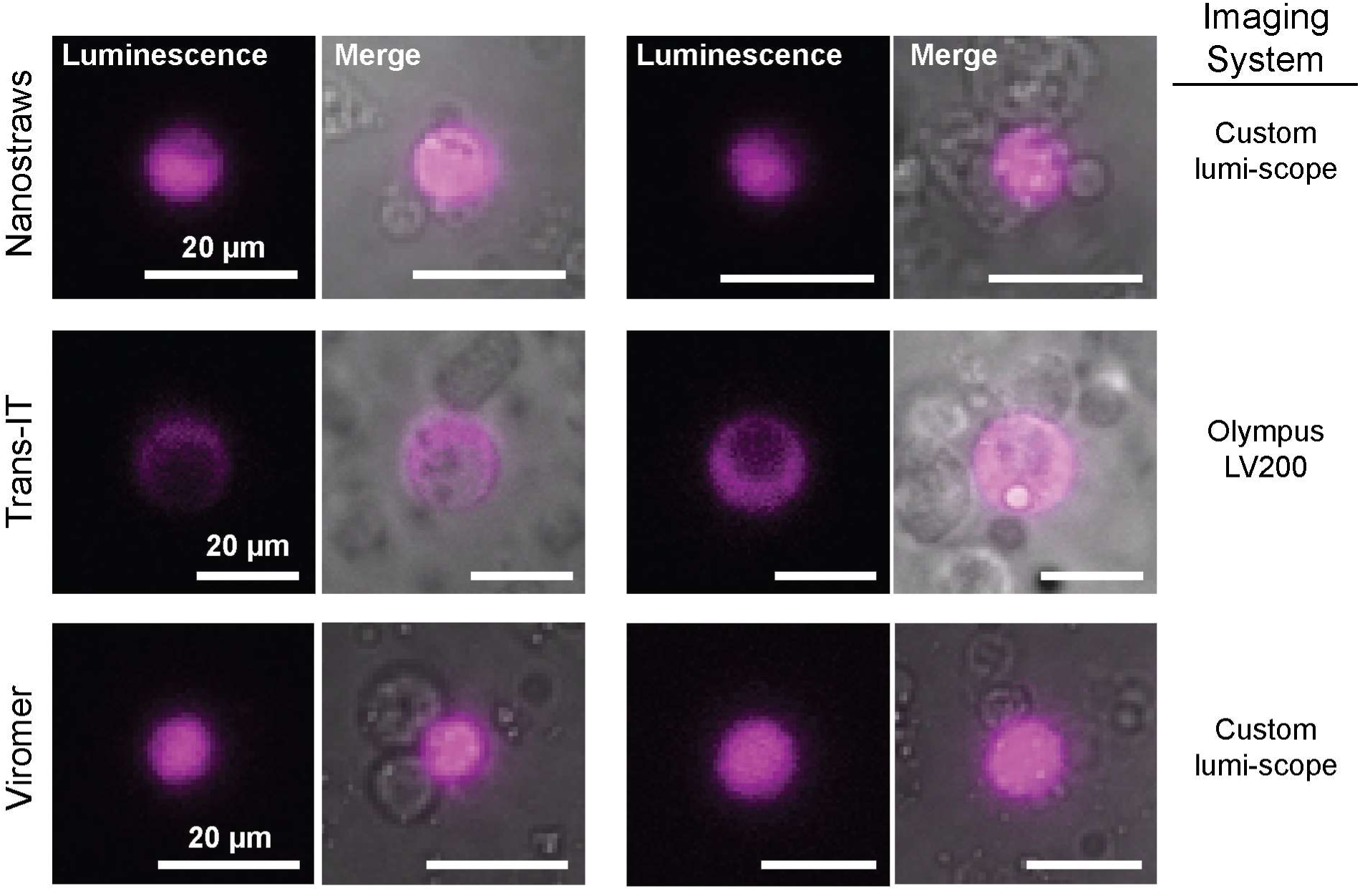
Luminescence imaging of transfected cells, related to Figure 2. Representative images of planarian cells transfected with either nanostraws (300 nm × 200 nm, 35V, 200 µs, 40 Hz) delivering RPL15-sNluc2 mRNA, Trans-IT delivering YB1-sNluc1 mRNA, or Viromer delivering RPL15-sNluc2, utilizing optimized transfection conditions for each method. Cells are imaged at 20 °C on either an Olympus LV200 luminescence microscope with a 100× oil immersion objective (Olympus: UPLXAPO100XO) or the custom-build luminescence microscope with a 100× oil immersion objective (Carl Zeiss: 1084-514) to observe cellular localization of luminescence (magenta).

**Figure S5:**
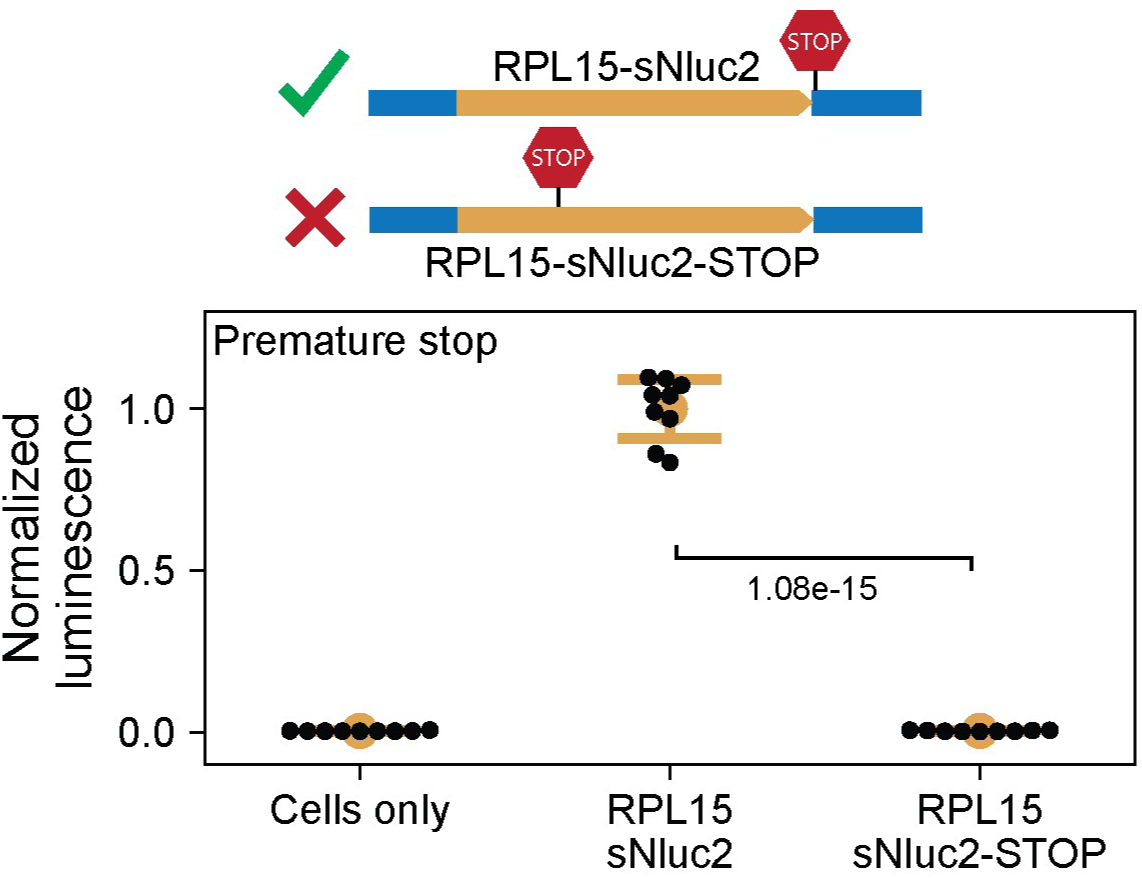
Premature stop codon abolishes Nluc expression, related to Figure 2. Luminescence from 200,000 dissociated planarian cells transfected with Viromer and RPL15-sNluc2 mRNA containing either a stop codon at the end of the 516 amino acid sequence or a variant containing a premature stop codon at residue 16. Luminescence is assayed 24 hpt. Statistics. All data points presented are technical replicates (n = 3) from independent biological replicates (n = 3) using mRNA produced from independent *in vitro* synthesis reactions. Luminescence measurements are normalized to the mean luminescence of RPL15-sNluc2. Error bars: standard deviation. ANOVA: p = 1.22e-18. p-value reported in the figure is calculated from a two-sided Welch’s t-test.

**Figure S6:**
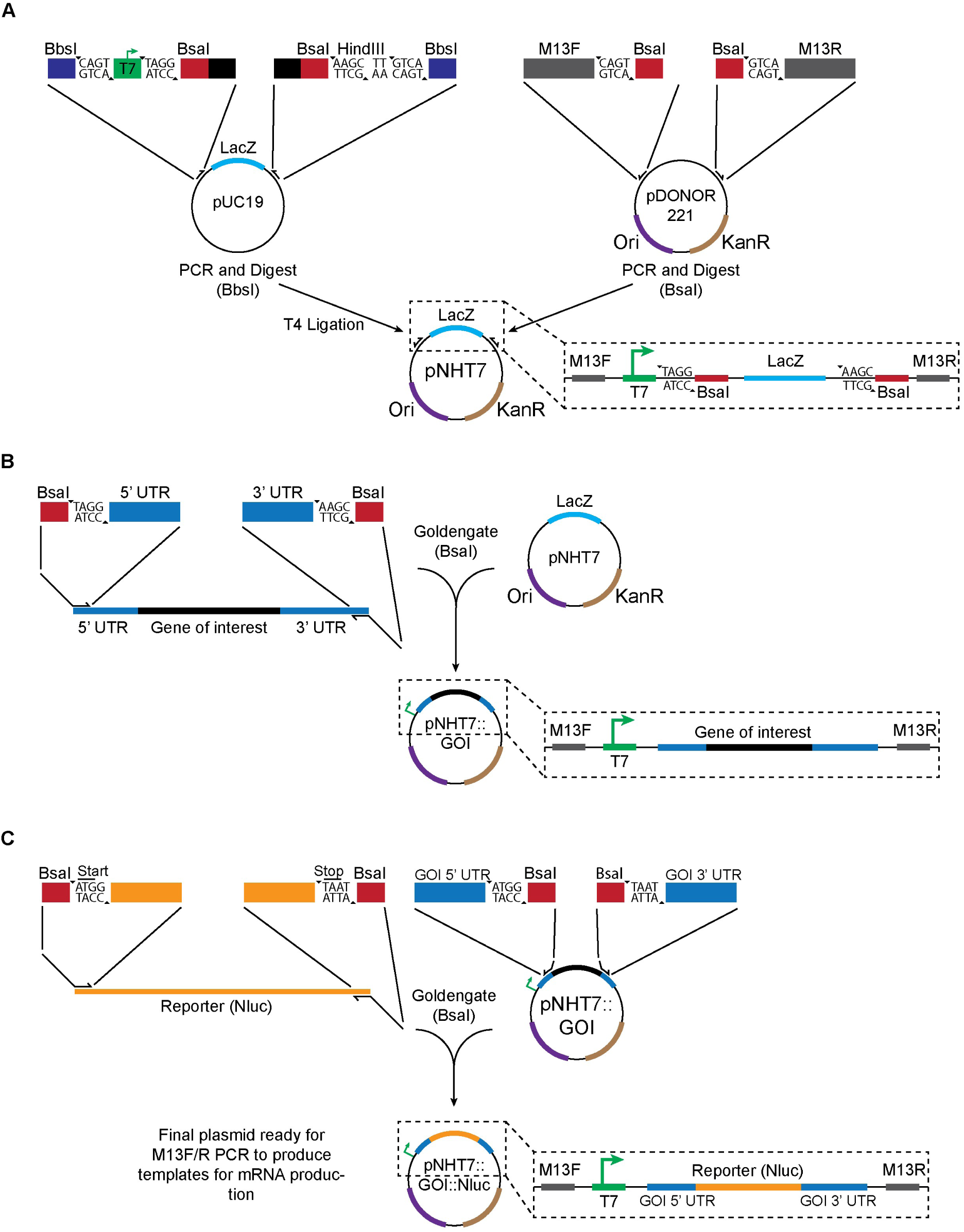
Cloning methodology, related to Figure 3. **(A)** A diagram showing the steps of generating pNHT7, the base vector used to clone endogenous planarian genes. The overall design is a LacZ cassette, derived from pUC19, flanked by BsaI restriction sites, which allow for subcloning of planarian genes into pNHT7. Upstream is a T7 promoter, which allows for IVT, and all of these are flanked by M13 forward and reverse primers, which are used to generate linearized templates for IVT through PCR. The backbone for pNHT7 is derived from the backbone from pDONOR221. **(B)** Isolating the 5’ and 3’ UTRs of a GOI from cDNA uses primers containing pNHT7 compatible BsaI restriction sites followed by 5’ and 3’ UTR sequences of the GOI. After amplification, the fragment containing the GOI can be cloned into pNHT7 though a standard golden gate reaction. **(C)** The Nluc reporter is inserted between the UTR sequences by using primers containing BsaI restriction sites to prime the 5’ and 3’ ends of the reporter as well as outward facing primers which add complementary BsaI restriction sites and prime the end of the 5’ UTR and beginning of the 3’ UTR for the gene captured in pNHT7. These two amplicons can then be purified and assembled via a golden gate reaction. All primer sequences are provided in **Table S4**.

**Figure S7:**
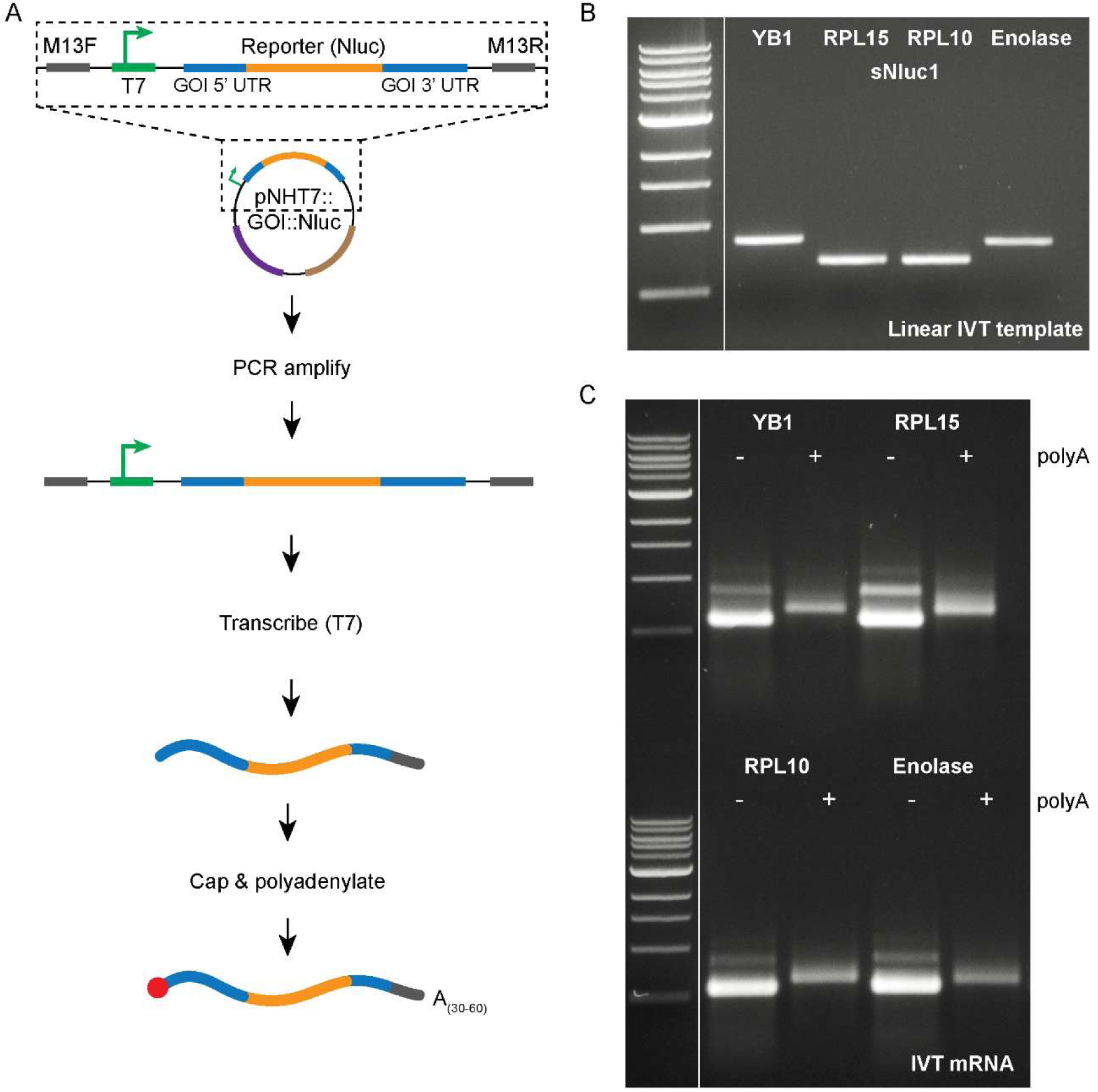
Expected *in vitro* transcription results, related to Figure 3. **(A)** Diagram of IVT. **(B)** EtBr stained 1% agarose gel showing the linear amplification products. **(C)** Gel images showing the mRNA product before and after polyadenylation. Upon successful polyadenylation, the mRNA product should be shifted in size. Samples should be free of excessive degradation products or additional unexpected bands, which we find critical for successful transfections. The ladder used is the NEB 1 kb DNA ladder.

**Figure S8:**
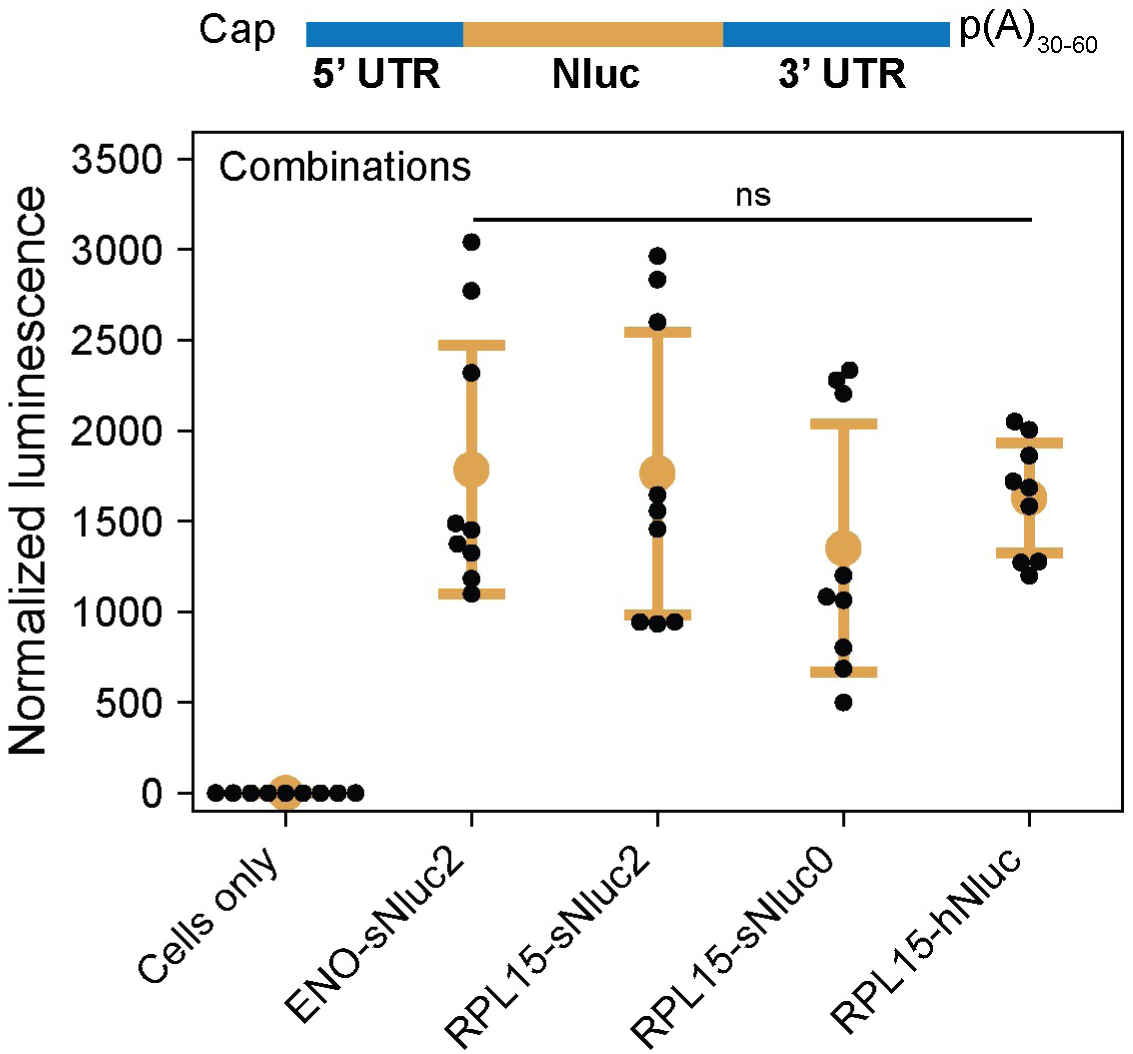
Combinations of UTR sequences with codon optimized Nluc variants, related to Figure 3. Luminescence from transfections using constructs that combine various UTRs with Nluc codon variants. All transfections are performed with 200,000 cells, from whole dissociated planarians, using 0.8 µL Viromer and 1 µg mRNA per well and assayed 24 hpt. Statistics. All data points presented are technical replicates (n = 3) from independent biological replicates (n = 3) utilizing mRNA produced from independent IVT reactions. Luminescence measurements are normalized to the mean value of the negative control (cells only). Error bars: standard deviation. Experimental conditions show no statistical significance (ANOVA: p = 0.51).

**Figure S9:**
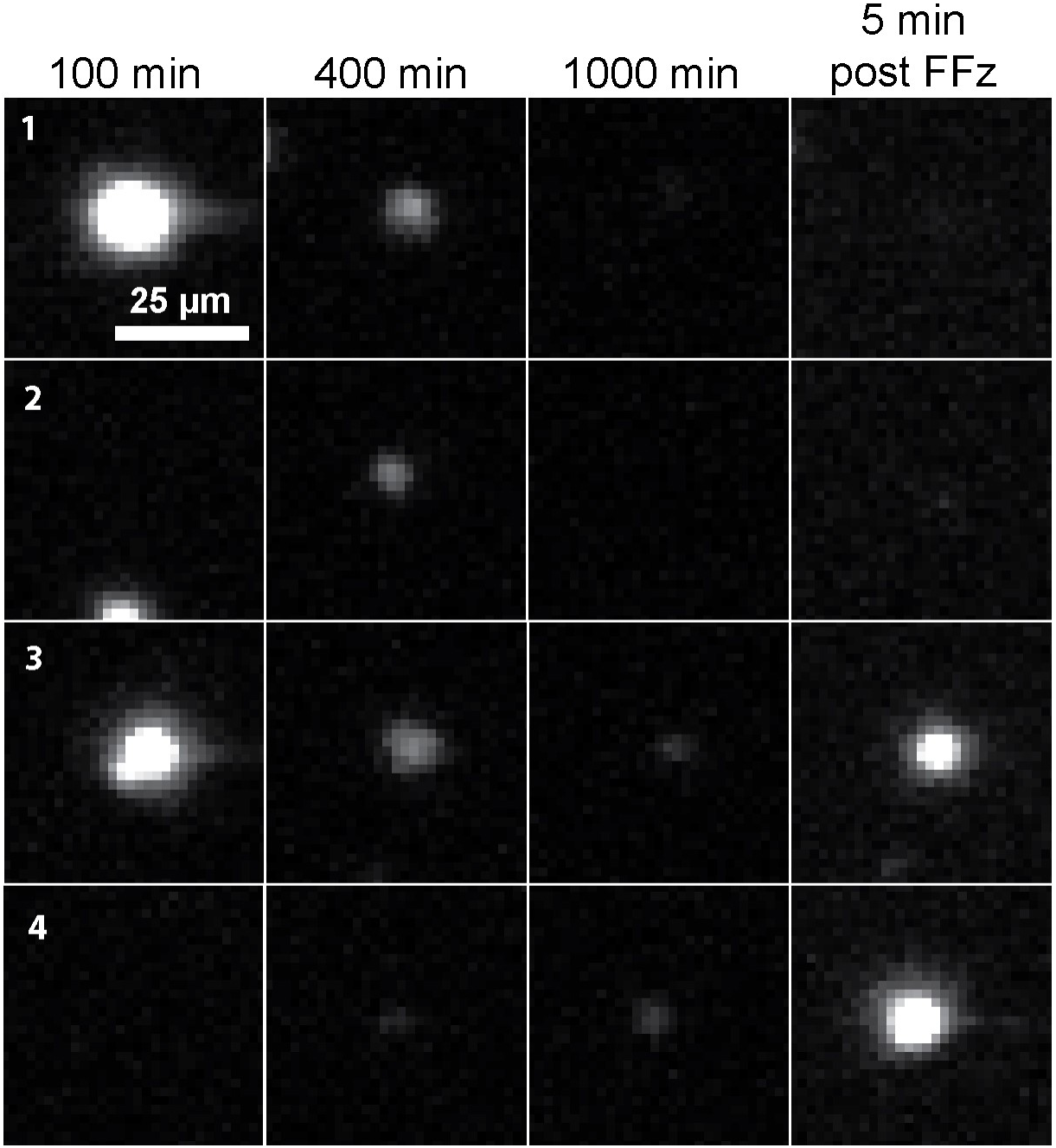
Cell responses to addition of fresh substrate, FFz, after 16 hr of live imaging, related to Figure 4. 200,000 dissociated planarian cells are transfected with 0.8 µL of Viromer and 1 µg of RPL15-sNluc2 mRNA and imaged through a 20× air objective (BoliOptics: BM03023431) at 1 frame per min with a 30 s exposure time for ∼17 hr on the custom-built luminescence microscope. After the first imaging session, we add 25 µL of 1:20 FFz diluted in Iso-L15, allow the cells to incubate for 5 min, and acquire an additional post-addition image. Cells exhibit four types of responses. 1. An early-transfected bright cell which does not respond to fresh FFz. 2. A late-transfected dim cell which does not respond to fresh FFz. 3. An early-transfected bright cell which does respond to fresh FFz. 4. A late transfected dim cell which becomes substantially brighter upon addition of fresh FFz.

**Figure S10:**
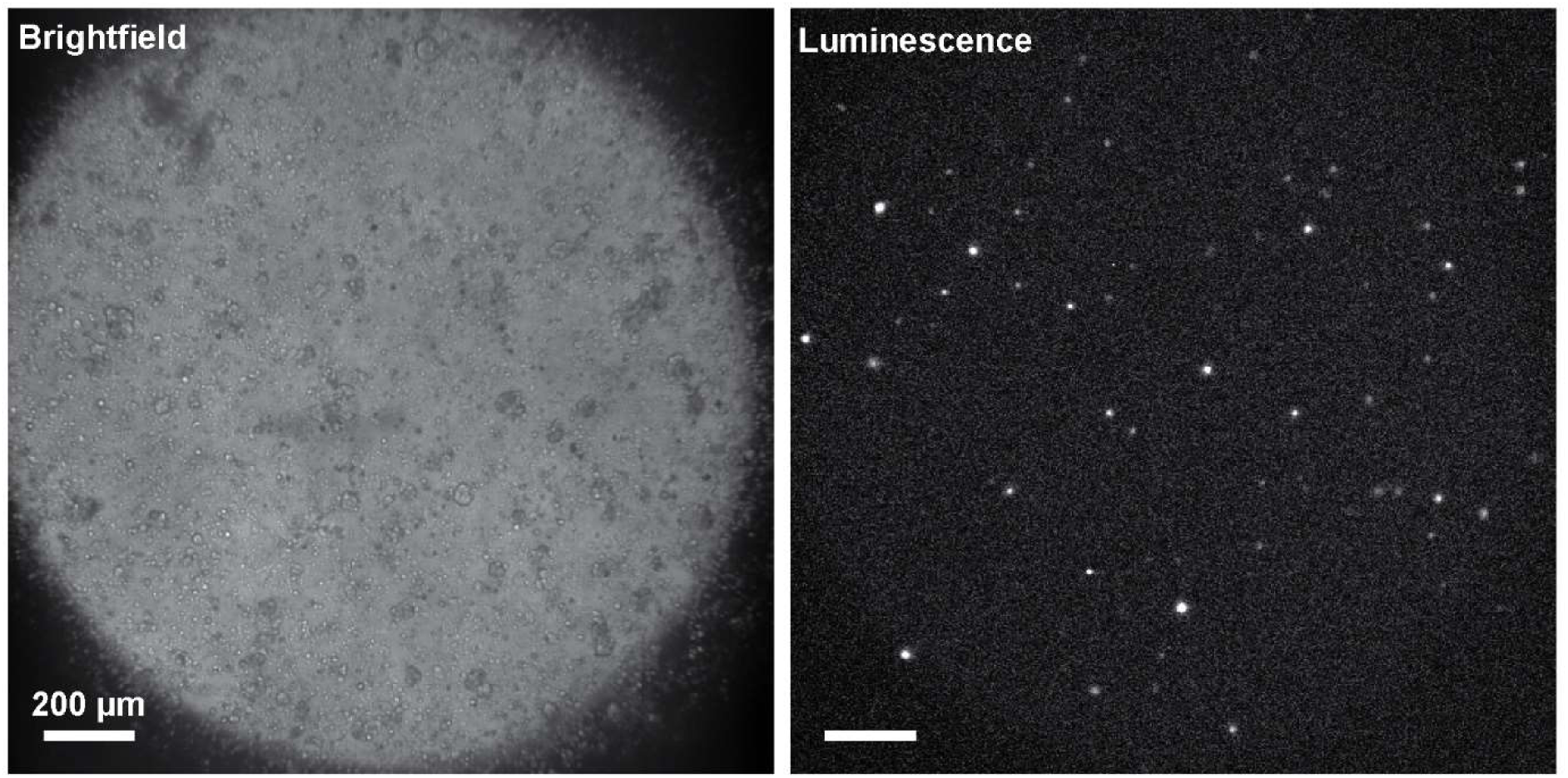
Imaging of dissociated cells after *in vivo* transfection, related to Figure 6. At 4 hpt using the method described in Figure 6, 10 animals are amputated posterior to the pharynx but anterior to the injection site. The tails are pooled and dissociated, then 1/3 of the cells are plated on a Concanavalin A-treated glass-bottom dish (35 mm). The cells are allowed to settle and adhere for 1 hr, then FFz is added (1:250), and the cells are imaged with a 20× air objective (BoliOptics: BM03023431), with an exposure time of 30 s, and a gain of 500.

**Figure S11:**
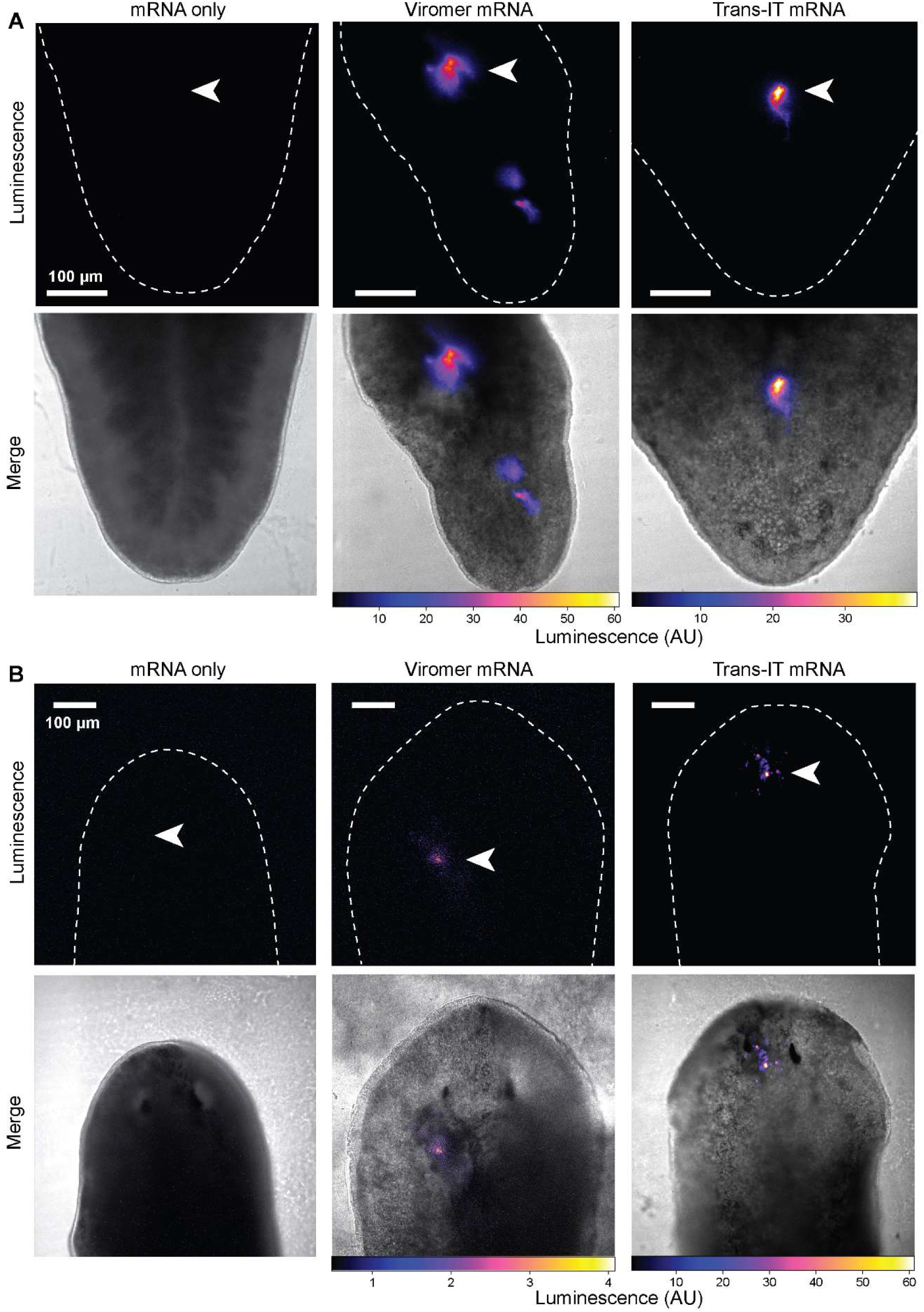
Live imaging of transfected animals, related to Figure 6. **(A)** Planarians injected along the tail midline. **(B)** Planarians injected at the anterior near the left eye cup. In both experiments live planarians are imaged on the Olympus LV200 luminescence microscope through a 20× air objective (Olympus: UPLXAPO20X). Negative control planarians are injected with mRNA alone. Images are acquired at 12 hpt. Animals are anesthetized and incubated for 15 min in furimazine (NanoGlo-Live, Promega) supplemented with 1% DMSO to aid in permeabilization. Luminescence images are acquired using an exposure of 60 s. All experiments are performed using identical exposure time and digital gain. Transfections are performed according to the conditions used in Figure 6B-C. Dashed lines: tail boundary; arrows: injection sites.

**Figure S12:**
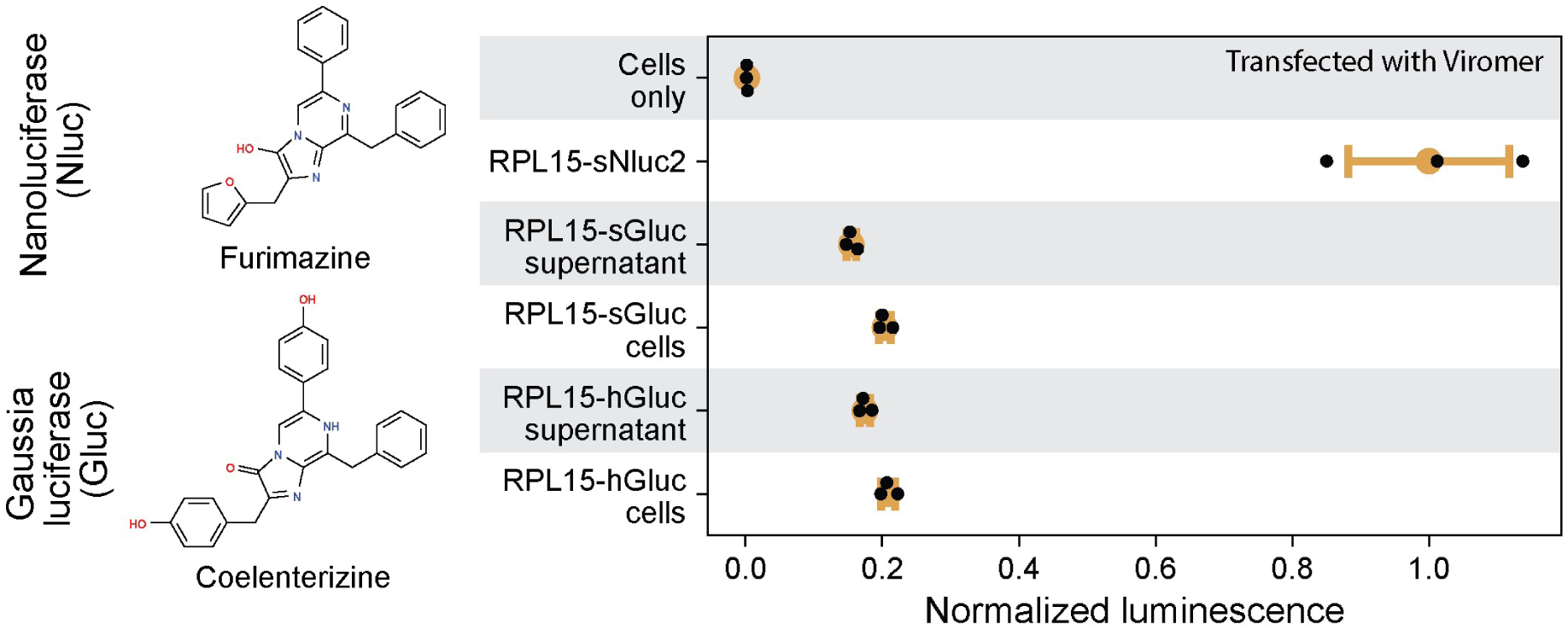
Expression of Gluc in planarian cells. 200,000 bulk dissociated cells are transfected with either RPL15-sGluc (planarian codon optimized), or RPL15-hGluc (human codon optimized) mRNA *in vitro* using Viromer. The cellular pellet and supernatant are separated and assayed independently using the substrate coelenterazine (chemical structure shown on the left) at 24 hpt, as Gluc protein is often secreted (Badr et al., 2007). Indeed, we detect clear signal from both the supernatant and the pellet. Note that luminescence is lower in Gluc transfected cells compared to positive controls transfected with Nluc. Chemical structures of the Nluc and Gluc substrates are shown on the left. All data points represent technical replicates utilizing mRNA prepared from the same batch. Luminescence measurements are normalized to the mean value of RPL15-sNluc2. Error bars, standard deviation.

**Figure S13:**
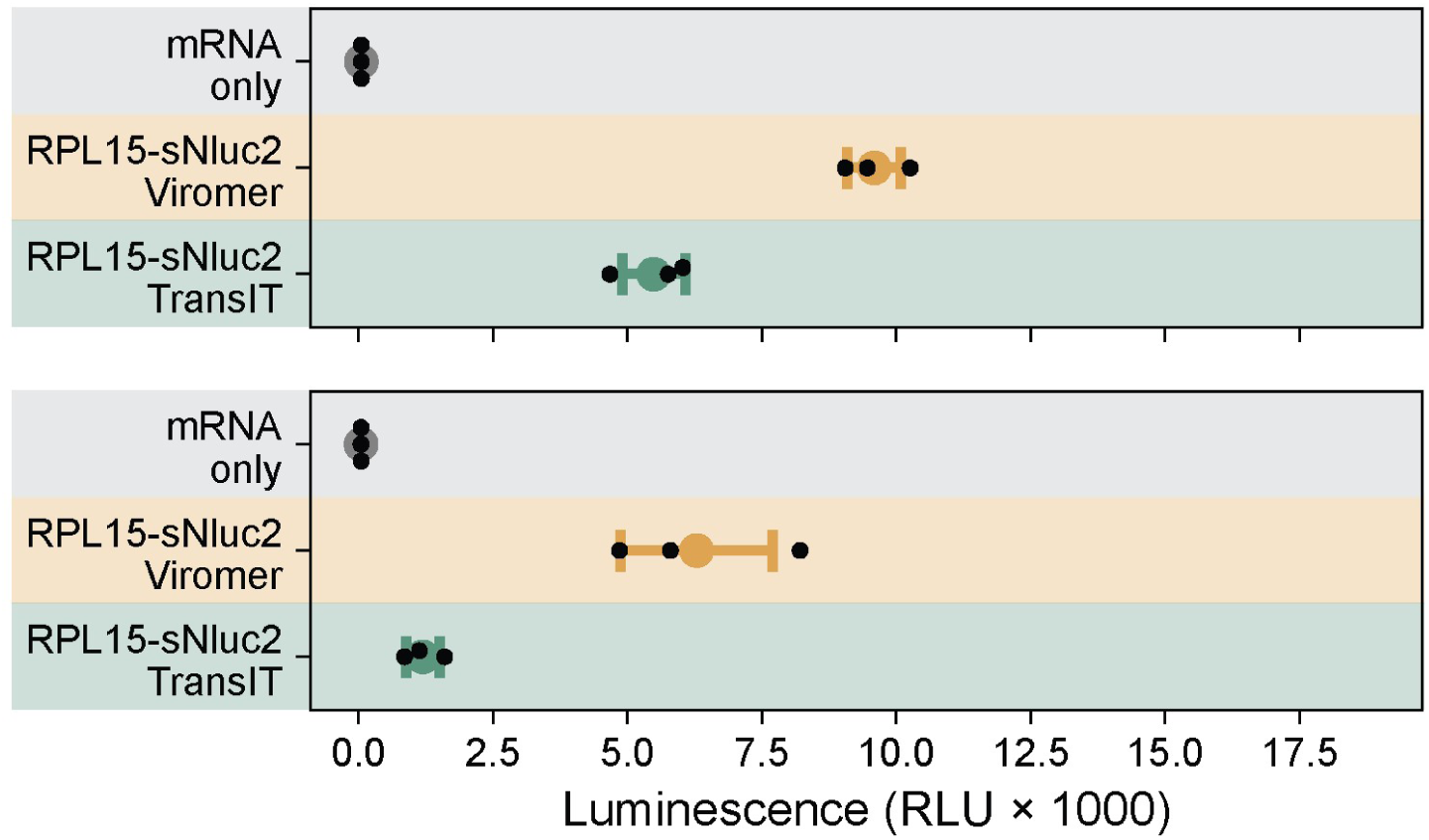
Nluc expression in *Schmidtea polychroa* cells. Transfections of 200,000 dissociated cells from *S. polychroa* either strain #200 (top) or strain #55 (bottom). Cells are transfected with either 0.8 µL Viromer (yellow) or 2 µL Trans-IT (green) and 1 µg of RPL15-sNluc2 mRNA. Luminescence is assayed 24 hpt. Data points represent technical replicates using mRNA prepared from the same batch. Error bars: standard deviation.

**Figure S14:**
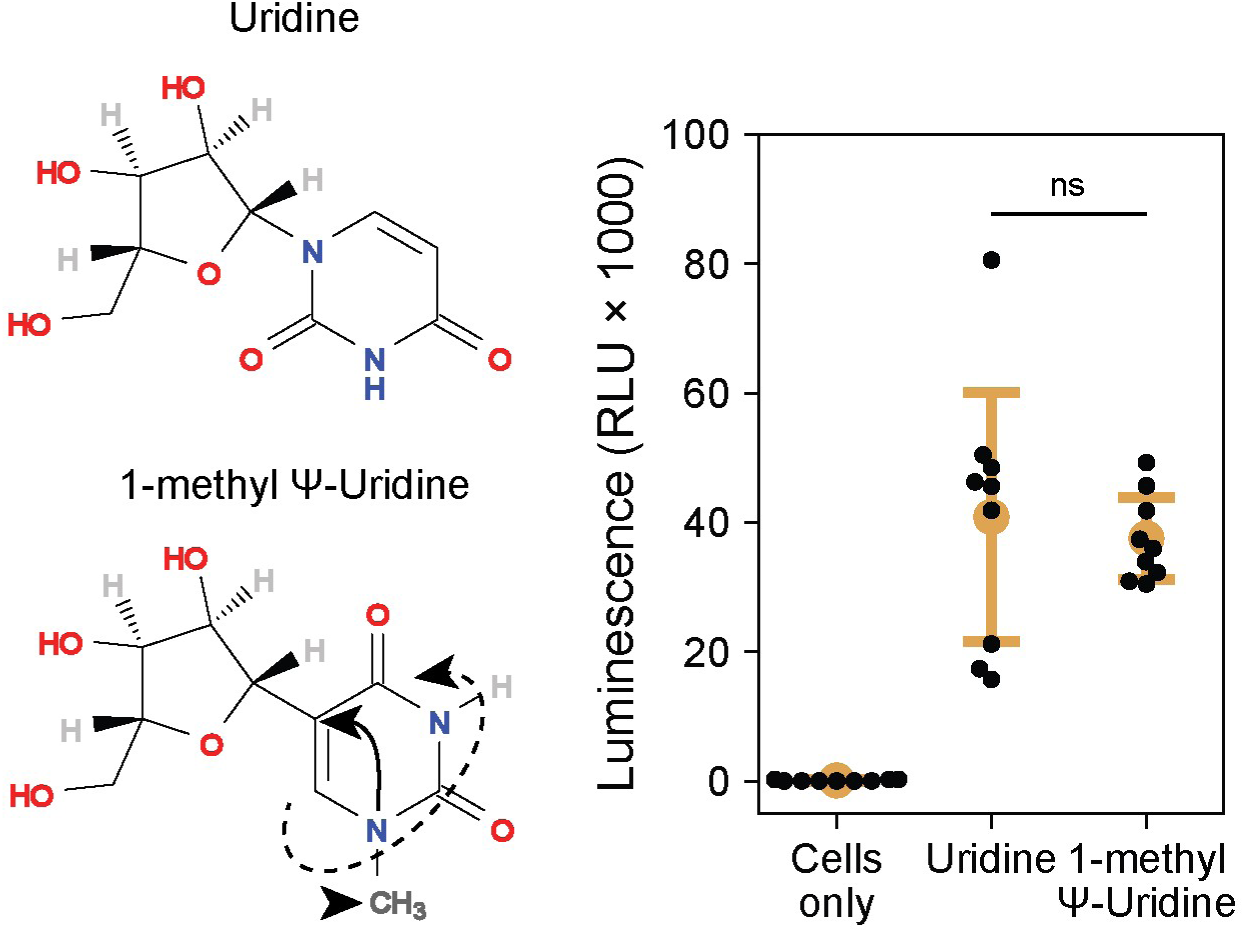
Assessment of m1Ψ on mRNA expression in planarian cells. Recent reports suggest that the base analog 1-methyl pseudouridine (m1Ψ) may increase translation, improve mRNA stability, and decrease immunogenicity (Andries et al., 2015; Svitkin et al., 2017). This modification is also currently being used in recently developed mRNA vaccines against SARS-CoV-2. We compare transfections with RPL15-sNluc2 mRNA containing either rUTP or m1Ψ (chemical structures shown on the left). We find both to be active, with no significant difference between the two samples (Welch’s t-test), suggesting that the observed benefits of m1Ψ modified mRNA in mammalian systems may not be directly transferrable to more basal invertebrates like planarians. Arrow: the uracil group is flipped 180 degrees and attached by a carbon-carbon bond rather than the usual carbon-nitrogen bond in pseudouridine. Arrowhead: m1Ψ contains an additional methyl group. All transfections are conducted with 0.8 µL of Viromer mRNA and 1 µg of mRNA. 200,000 cells are incubated at 20°C, and luminescence is measured at 24 hpt. All data points presented are technical replicates (n = 3) from independent biological replicates (n = 3) utilizing mRNA produced from independent IVT reactions. Error bars: standard deviation.

### Supplemental Tables

**Table S1: Nluc variants used in this study.** Coding sequences with different codon usage bias are listed with their respective GC content, and codon adaptation index (CAI) calculated from https://ppuigbo.me/programs/CAIcal/.

**Table S2: Conditions for screening transfection reagents.** The conditions follow the manufacturer’s protocol. Reagent 1 is the main transfection reagent provided with the kit (e.g., Lipofectamine 3000, Trans-IT mRNA, Viromer mRNA). Reagent 2 is a reagent provided by some but not all kits. For Trans-IT it is the Boost reagent, and for Lipofectamine 3000, it is the Transfection Enhancer reagent. **Diluting** Buffer is the solution used to dilute the mRNA and transfection reagents. Some kits provide a specific buffer to use (Viromer) while others require serum-free medium, which is L-15 in our experiments. Incubation is done in room temperature and is recommended to allow the reagent and mRNA to complex before adding to cells.

**Table S3: List of constructs used in this study.** Corresponding plasmid maps are provided in GenBank (.gb) format as **File S1**. All constructs are available through Addgene.

**Table S4: Primers used in this study.** Listed are primer ID, conventional name, a short description of the usage, and the sequence from 5’ to 3’.

### Supplemental Videos

**Video S1:** Time-lapse luminescence imaging of RPL15-sNluc2 expressing dissociated planarian cells immediately after transfection. This video corresponds to the data shown in **Figure 4D-H**.

**Video S2:** Example of luminescent traces for individually segmented cells taken from **Video S1**.

**Video S3:** Montage of luminescence imaging in live planarians, which are decapitated to reduce motility. This video corresponds to the data shown in **Figure 6E**.

**Video S4:** Live luminescence imaging in planarians immediately post injection. This video corresponds to the data shown in **Figure 6F-G**.

### Supplemental File

**File S1:** An archive containing maps for all plasmids (GenBank, .gb) generated and used in this work.

